# Functional analysis of a common BAG3 allele associated with protection from heart failure

**DOI:** 10.1101/2021.10.06.463213

**Authors:** Juan A Perez-Bermejo, Luke M Judge, Christina L Jensen, Kenneth Wu, Annie Truong, Jaclyn J Ho, Matthew Carter, Wendy V Runyon, Robyn M Kaake, Ernst Pulido, Hannah L Watry, Mohammad A Mandegar, Danielle L Swaney, Po-Lin So, Nevan J Krogan, Bruce R Conklin

**Affiliations:** Gladstone Institutes, San Francisco, CA, United States; UCSF Department of Pediatrics, San Francisco, CA, United States; Tenaya Therapeutics, South San Francisco, CA, United States; UCSF Quantitative Biosciences Institute (QBI), San Francisco, CA, United States; UCSF Department of Cellular Molecular Pharmacology, San Francisco, CA, United States; UCSF Department of Medicine, San Francisco, CA, United States; Innovative Genomics Institute, Berkeley, CA, United States

## Abstract

Multiple genetic association studies have correlated a common allelic block linked to the BAG3 gene with a decreased incidence of heart failure, but the molecular mechanism for such protection remains elusive. One of the variants in this allele block is coding, changing cysteine to arginine at position 151 of BAG3 (rs2234962-BAG3^C151R^). Here, we use induced pluripotent stem cells (iPSC) to test if the BAG3^C151R^ variant alters protein and cellular function in human cardiac myocytes. Quantitative protein interaction network analysis identified specific changes in BAG3^C151R^ protein interaction partners in cardiomyocytes but not in iPSCs or an immortalized cell line. Knockdown of BAG3 interacting factors in cardiomyocytes followed by myofibrillar analysis revealed that BAG3^C151R^ associates more strongly with proteins involved in the maintenance of myofibrillar integrity. Finally, we demonstrate that cardiomyocytes expressing the BAG3^C151R^ variant have improved response to proteotoxic stress in an allele dose-dependent manner. This study suggests that the BAG3^C151R^ variant increases cardiomyocyte protection from stress by enhancing the recruitment of factors critical to the maintenance of myofibril integrity, hinting that this variant could be responsible for the cardioprotective effect of the haplotype block. By revealing specific changes in preferential binding partners of the BAG3^C151R^ protein variant, we also identify potential targets for the development of novel cardioprotective therapies.

## Introduction

Heart failure is a major cause of mortality, with increasing incidence each year^1^. One of the main causes of non-ischemic heart failure is dilated cardiomyopathy (DCM), which has a strong genetic component^1–3^. Although many genetic association studies have identified risk loci associated with DCM, the translation of these discoveries into therapeutic strategies remains challenging. This is partly due to a lack of mechanistic insights at the molecular level, making it difficult to progress from correlation to causality. In addition, most of the genetic variants identified in genetic association studies are rare and disease-causing, so the mechanistic information is applicable to only a small subset of patients. On the contrary, variants associated with a decreased risk of disease (protective) have the potential to be utilized for the development of therapeutic strategies that are applicable to a much broader range of subjects^4^. Recently, multiple large genome wide association studies have implicated a common haplotype block, which overlaps with the BAG3 (Bcl-2 associated athanogene 3) gene, with an apparent decreased incidence of DCM or with improved cardiac ejection fraction^5–13^ (for a summary of these studies, see ***Table 1***). Despite the strong genetic association, the molecular basis of the apparent protective effect remains unknown. A mechanistic understanding of this protective allele could aid in the development of novel cardioprotective therapies.

**Table 1.**
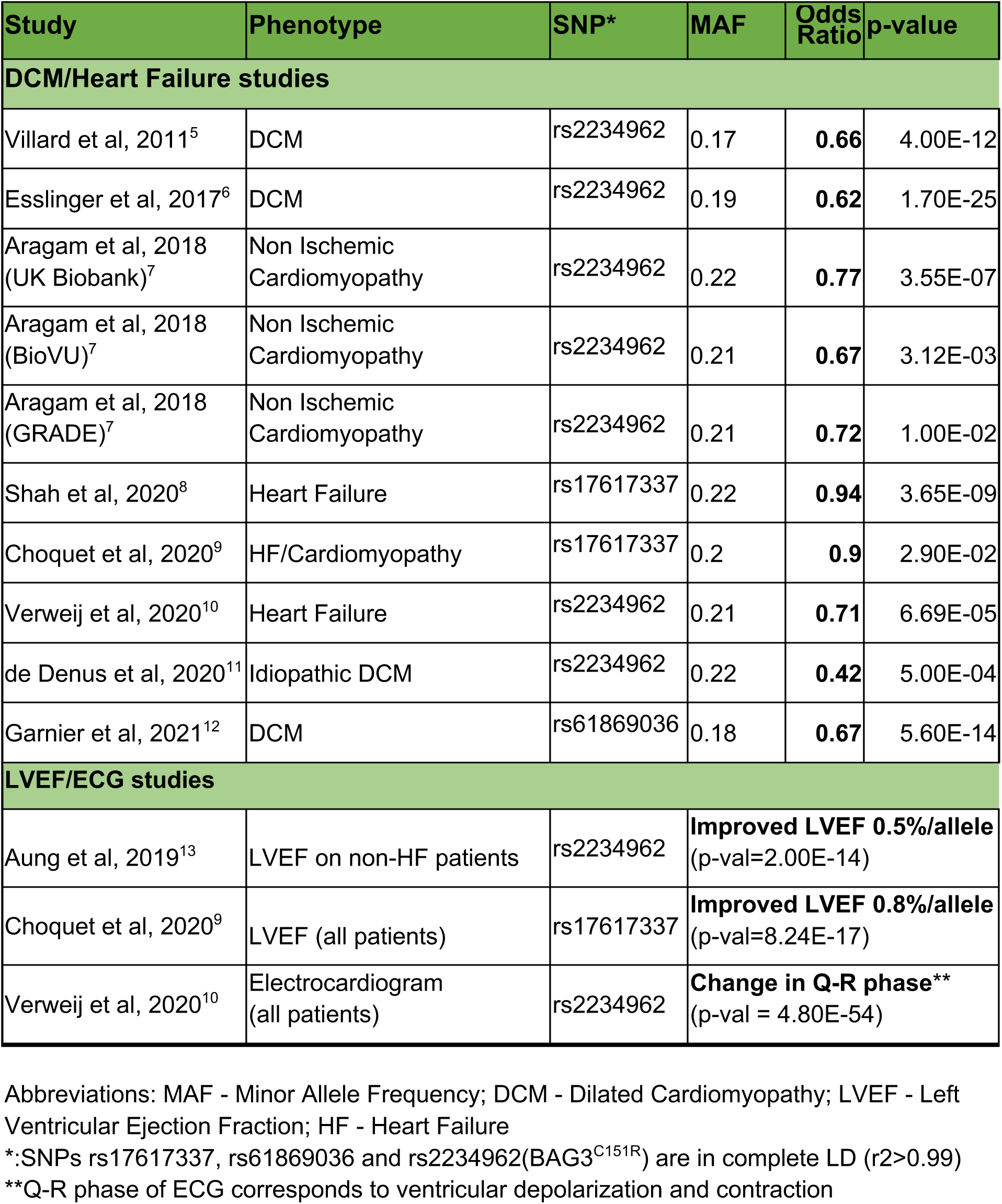
Summary of genome wide association studies supporting the putative cardioprotective effect of the BAG3 haplotype block.

A combination of clinical case reports and large genetic association studies have implicated BAG3 loss-of-function variants with DCM^5, 7, 14–20^. BAG3 was first described as an HSP70 co-chaperone, while successive studies have found it to be a multifunctional hub for a diverse set of cellular processes, mostly centered around protein quality control and selective autophagy^21–23^. In cardiac and skeletal muscle, a functioning protein quality control network is essential to dynamically maintain myofibrillar structures through constant cycles of contraction^24^. Accordingly, BAG3 impairment results in heart and skeletal muscle pathology^25–28^. These studies indicate the BAG3 protein has an essential role in muscle, particularly in cardiomyocytes, which lack replicative potential. In particular, BAG3 binding to the HSP70 chaperones is essential for cardiac health, as indicated by multiple pathogenic mutations that reduce this interaction^14, 15, 17, 18, 29^. Interestingly, the cardioprotective BAG3 allele block contains one coding variant, rs2234962, which leads to the BAG3^C151R^ amino acid transition. However, the C151R change is located distant from the HSP70 binding domain, in an intrinsically disordered region with no known function. Based on these observations, we hypothesize that the BAG3^C151R^ could induce a novel functional effect in the BAG3 protein and be responsible for the effect of the cardioprotective haplotype block.

Recent studies have demonstrated that induced pluripotent stem cell (iPSC) modeling can be used to elucidate the functional effects of disease risk variants identified in genetic association studies^30–33^. Here, we have leveraged the power of iPSC technology and genome editing to examine the impact of the C151R aminoacid change in the function of the BAG3 protein, and on the response of iPSC-derived cardiomyocytes to stress. Our findings confirm that the BAG3^C151R^ variant has a protective effect *in vitro*, and could be responsible for the effect of the cardioprotective haplotype block. We also discovered BAG3^C151R^-associated alterations in protein interactions that are cardiomyocyte specific and hint at novel mechanisms for cardioprotection.

## Results

### BAG3^C151R^ quantitatively alters the profile of BAG3 interaction partners in cardiomyocytes

The BAG3 allele block associated with reduced incidence of heart failure comprises at least seven common variants in almost complete linkage disequilibrium across an ∼18Kb region, of which only the rs2234962 variant is exonic ***(Fig 1A; Fig S1A)***. The block presents with an allele frequency of roughly 20% in European and South Asian populations ***(Fig 1B; Fig S1B)***. The rs2234962 variant results in the BAG3^C151R^ amino acid change, which is located in a region of the protein with no annotated domain or function ***(Fig 1C; Fig S1C-D)***. Since BAG3 is a scaffold protein with a wide range of binding partners, we reasoned that if this variant indeed alters BAG3 function, it would likely do so via alterations in the profile of protein interactions. Presumably, these changes would be different from those caused by a pathogenic variant such as the BAG3^E455K^, which lies in a highly conserved region of the protein ***(Fig S1E)*** and has been shown to disrupt protein interactions in the mouse heart^20, 27^***(Fig 1C)***.

**Figure 1.**
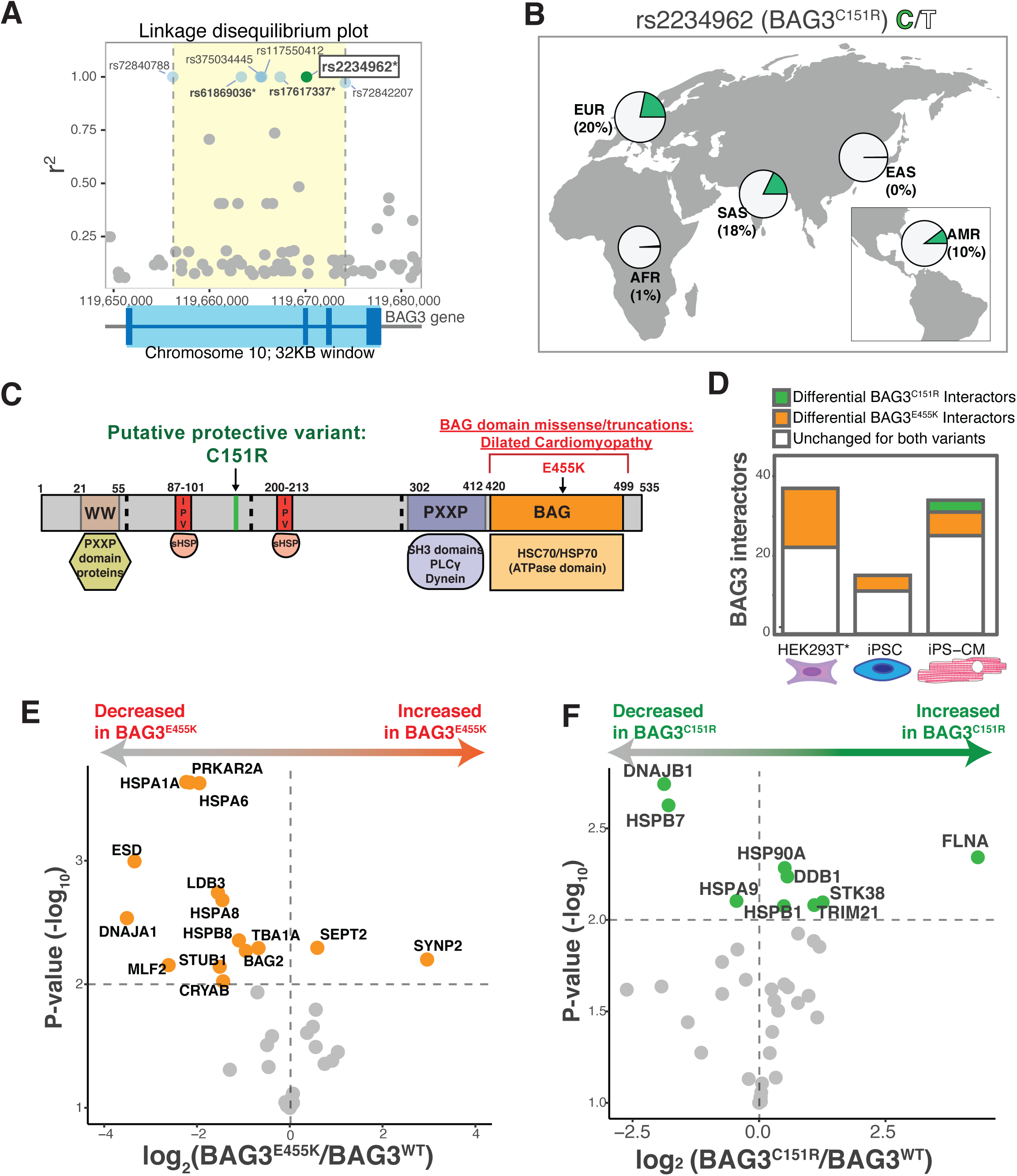
The putative cardioprotective variant rs2234962-BAG3^C151R^ results in a change in the profile of BAG3 co-precipitation partners in a cardiomyocyte background. (A) Top: linkage disequilibrium plot centered around the rs2234962 variant (green) showing six other genetic variants (blue) in complete disequilibrium that form a haplotype block. *: Variants used in genetic association studies. Data for Central European population (CEU). Bottom: gene structure of the overlapping BAG3 gene (light blue: introns, dark blue: exons). (B) Allele frequencies across different super populations from the 1000 Genomes database. (AFR: African, AMR: Ad Mixed American, EAS: East Asian, EUR: European, SAS: South East Asian). (C) Schematic of BAG3 protein domain structure, highlighting some relevant interaction partners and indicating the location of the variants used in this study (arrows). Numbers indicate amino acid boundaries for annotated domains. Dashed lines indicate exon-exon boundaries. Areas of unknown function are colored in grey. (D-F) Interaction analyses for three different BAG3 variants (BAG3^WT^, BAG3^C151R^, BAG3^E455K^). (D) Bar plot of the number of differential interactors (co-precipitation partners that interacted significantly more or less with BAG3 in either variant compared to BAG3^WT^) identified using APMS in three different cell backgrounds - HEK293T cells (*: indicates BAG3 variant overexpression), undifferentiated human iPSC, and differentiated iPS-CM. Significantly different binding partners of BAG3^C151R^ were only identified in the iPS-CM background. *N=4.* (E) Volcano plot depicting the intensity ratio of co-precipitation of each interactor with BAG3^E455K-3xFLAG^ relative to BAG3^WT-3xFLAG^ in iPS-CM. (F) Volcano plot depicting the intensity ratio of co-precipitation of each interactor with BAG3^C151R-3xFLAG^ relative to BAG3^WT-3xFLAG^ in iPS-CMs. Horizontal dashed line indicates statistical significance threshold (adjusted p-value <0.01) and vertical dashed line indicates no ratio change. BAG3^C151R^ results in a significant change of interactions that are different from those affected by the pathogenic variant. *N*=4.

To test this hypothesis, we first performed affinity purification coupled to mass spectrometry (APMS) to characterize the stable protein interaction partners of a set of BAG3 variants overexpressed in a non-muscle immortalized cell line, HEK293T. We observed a significant loss of co-precipitation partners (mostly HSP70 family members) for the pathogenic BAG3 variants ***(Fig 1D; Fig S2; Table S1)***. This result validated our approach, however no changes in protein interactions were observed for BAG3^C151R^. We postulated that to see a difference for this variant, performing these experiments in a more relevant cell type, the cardiomyocyte, was necessary. We also reasoned that since BAG3 protein levels are tightly regulated, it would be important to use endogenous levels of expression to study interaction partners. We therefore generated a series of isogenic induced pluripotent stem cell (iPSC) lines bearing the BAG3^C151R^ variant, the pathogenic variant BAG3^E455K^, or the non-disease associated wild-type variant (BAG3^WT^), with a 3xFLAG epitope tag fused to the endogenous BAG3 gene ***(Fig S3)***. The cell lines were differentiated into cardiomyocytes (iPS-CMs) and we used APMS to analyze the BAG3 protein complexes in both iPSCs and iPS-CMs. As expected, we observed a difference in the significant stable partners of the BAG3^E455K- 3xFLAG^ variant compared to BAG3^WT-3xFLAG^ in both iPSCs and IPS-CMs ***(Fig 1D)***. In comparison to the HEK293T APMS results, we were also able to characterize both increasing and decreasing protein interactions for the BAG3^C151R-3xFLAG^ variant, but only in the iPS-CM background and not in undifferentiated iPSCs ***(Fig 1D).*** This highlights the cell type specificity of the BAG3^C151R^ variant function. In addition, we performed this APMS analysis for BAG3^WT-3xFLAG^ interactors in iPS-CMs under proteotoxic stress (using proteasome inhibitor bortezomib), and in an engineered inducible cell line overexpressing BAG3^WT-3xFLAG^***(Fig S4)***. This comprehensive characterization allowed us to obtain a refined list of stable BAG3 interactors in a cardiomyocyte background ***(Fig S5)***. This list included HSP70 chaperones and co-chaperones as well as other partners such as small heat shock proteins, sarcomeric factors and actin-associated proteins. It also showed a large difference in interactors identified when BAG3 was overexpressed, highlighting the importance of performing functional analyses with proteins expressed at endogenous levels ***(Fig S5D)***.

To quantify the changes in co-precipitation efficiencies of BAG3 interaction partners across variants, we performed a quantitative targeted proteomics analysis on the iPS-CM APMS samples. While the pathogenic variant BAG3^E455K-3xFLAG^ resulted in a generalized loss of protein interactions compared to BAG3^WT^ (mostly HSP70 chaperones and co-chaperones) ***(Fig 1E)***, the BAG3^C151R-3xFLAG^ variant displayed a significant change in a different subset of binding partners (**Fig 1F**). In particular, there was an increase in the interaction with actin-binding protein Filamin A, hippo pathway kinase STK38, and E3 ubiquitin protein ligases DDB1 and TRIM21, while interaction with small heat shock protein HSPB7 and co-chaperone DNAJB1 were decreased compared to BAG3^WT^.

Taken together, these results suggest that the putative protective variant BAG3^C151R^ influences BAG3 protein function by modifying its cardiomyocyte-specific profile of protein interactions.

### BAG3^C151R^ engages factors involved in the maintenance of sarcomeric homeostasis

Many stable BAG3 interaction partners identified here (such as those that significantly changed in the BAG3^C151R^ background) have no known roles in cardiomyocyte function, particularly in the context of BAG3 mediated protein homeostasis. Cardiomyocytes are contractile cells and the structure and integrity of their myofibrils is closely associated to their functional output and pathological status. Correspondingly, myofibrillar disarray is a hallmark of BAG3-related disease^19, 25, 34^. We therefore hypothesized that some BAG3 partners might also play a role in the maintenance of sarcomeric integrity in iPS-CMs. To test this hypothesis in an unbiased manner, and with a focus on BAG3-specific features, we set up a pipeline for systematic iPS-CM gene knockdown followed by unbiased sarcomere structure analysis of microscopy images ***(Fig 2A)***. Images from different gene knockdowns were scored using a supervised learning classifier that ranked sarcomere staining images by similarity to a BAG3 knockdown phenotype or a control phenotype, generating a ‘BAG3 sarcomere score’. Visual inspection of these two training conditions revealed that iPS-CMs with a BAG3 knockdown were more likely to collapse (with completely aggregated myofibrils) and generally presented a lower myofibril density accompanied with increased sarcomere disarray and staining gaps ***(Fig 2B; Fig S6A-C)***. Our scoring scheme accurately classified BAG3 knockdown and control test images using only sarcomere staining images as input, and the obtained score correlated with BAG3 protein levels ***(Fig 2C, Fig S7A-C)***.

**Figure 2.**
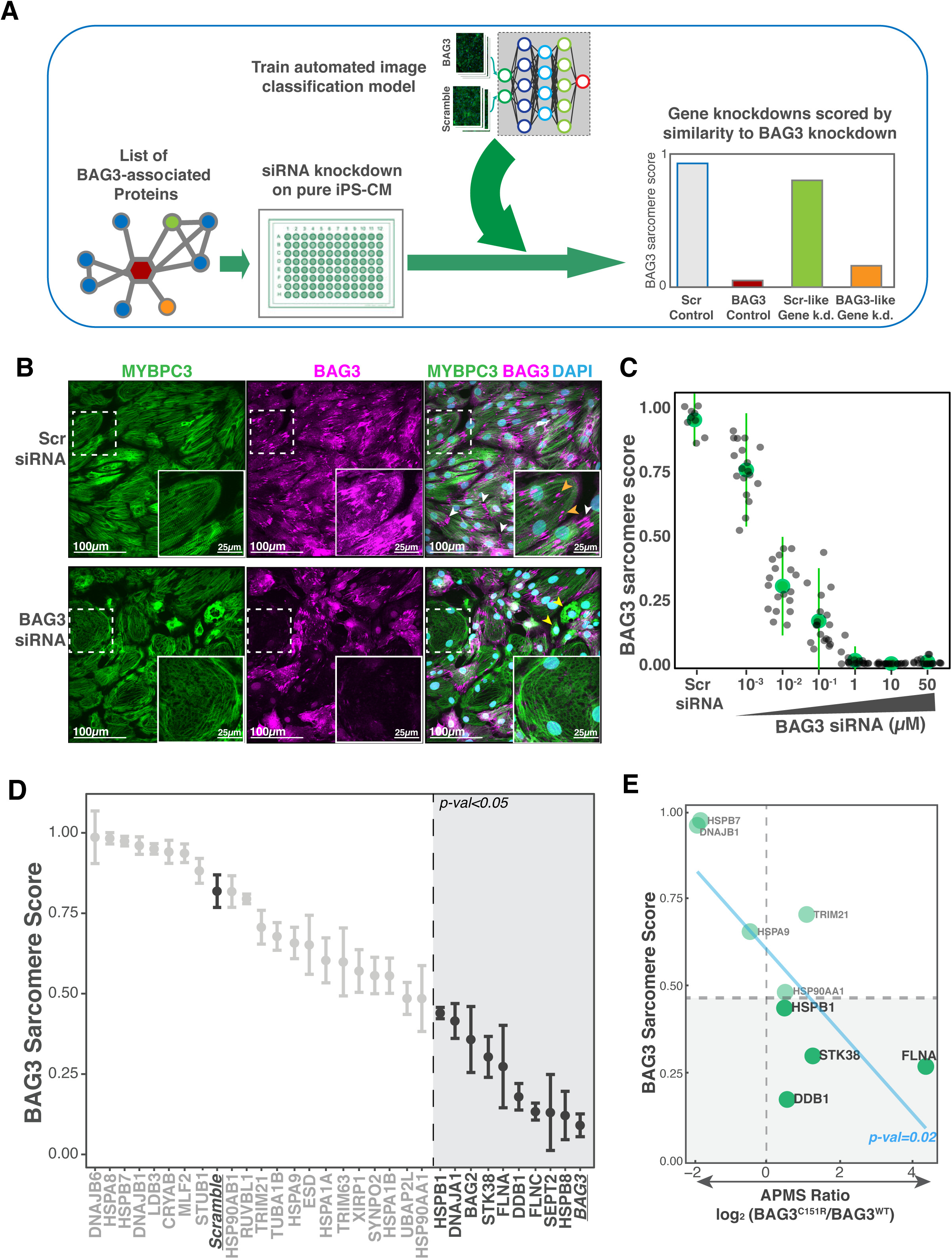
BAG3^C151R^ favors binding to factors required for maintenance of myofibrillar integrity. (A) Diagram of the pipeline followed for automated analysis of myofibrillar staining images in iPS-CM. First, BAG3 associated genes are knocked down using siRNA. Then those images are put through a supervised learning scoring scheme trained on myofibrillar staining images of iPS-CM treated with Scramble (control) or BAG3 siRNA. Scores are closer to 1 if the image resembles a scramble condition and closer to 0 if similar to a BAG3 knockdown, as shown in the mock graph in A. (B) Representative images of iPS-CMs treated with control or BAG3 siRNA. BAG3 staining localized to the sarcomeric z-disk region and with high intensity in the MYBPC3 staining gaps (presumably myofibrillar breaks; orange arrowheads) and, in some cases, to the polar ends of cardiomyocytes (white arrowheads). BAG3 knockdown cardiomyocytes display a reduced density of myofibrils, which are also more disorganized, and more cells with collapsed/aggregated sarcomere staining (yellow arrowheads). Cells with the lowest BAG3 staining displayed a particularly disorganized sarcomere structure, with numerous gaps in MYBPC3 staining (see insert). (C) Image score correlate anticorrelated with the amount of BAG3 siRNA used. *N=* 9 images for Scr siRNA condition, *N*=18 for the rest. (D) BAG3 sarcomere score for all the BAG3 co-precipitation partners identified in this study. Internal controls Scramble and BAG3 are underlined. Vertical dashed line marks the threshold beyond which a knockdown scored significantly lower than the internal Scr control (no condition scored significantly higher than Scramble). Dots represent mean of 3 replicates from separate wells, each being the median score of 9 images from the same well. Error bars: SEM. P-val cutoff: 0.05 using a one-way ANOVA with post-hoc Dunnett test. (E) Knocking down proteins that interact stronger with BAG3^C151R^ than BAG3^WT^ phenocopies BAG3 insufficiency. Plot depicting the relationship between the sarcomeric score of a gene knockdown and the corresponding protein’s BAG3^C151R^/BAG3^WT^ co-precipitation intensity ratio. P-value obtained fitting a linear model. Pearson’s product-moment correlation: -0.75.

We then applied this phenotypic analysis pipeline to the knockdowns of all the protein interactors identified in our BAG3 interactome in addition to some other high-confidence BAG3-related proteins ***(Table S2)***. We found that knockdown of 9 of the 32 genes encoding BAG3 iPS-CM high confidence protein interactions (and 6 out of 19 genes from outside our APMS dataset) phenocopied BAG3 insufficiency **(Fig 2D, Fig S7D, Fig S8)**. None of these knockdowns significantly affected cardiomyocyte viability, and only one (HSPB8, a well-known BAG3 partner in protein quality control^26, 35^) significantly decreased BAG3 protein levels ***(Fig S7E-F)***. Interestingly, knockdown of FLNA, STK38, DDB1 and HSPB1, proteins that more strongly co-precipitated with BAG3^C151R-3xFLAG^ than BAG3^WT^, were among the phenotypes that most resembled BAG3 insufficiency. In fact, we observed an inverse relationship between the BAG3 sarcomere scores and the change in co-precipitation intensity for all BAG3^C151R-3xFLAG^ differential interactors ***(Fig 2E)***. This correlation was not observed for BAG3^E455K^ differential binding partners ***(Fig S7G)***.

Altogether, this data implicates a specific subset of BAG3 interactors as particularly important for the maintenance of myofibrillar integrity. It also suggests that BAG3^C151R^ variant has an increased ability to engage some of these specific factors, presumably improving their BAG3-associated role in the maintenance of the sarcomeric structure.

### BAG3-C151R protects against toxic proteasome inhibition in cardiomyocytes

Since BAG3 is a well-known stress responsive factor with a role in the maintenance of cell homeostasis under proteotoxic conditions, we next wanted to elucidate whether the BAG3^C151R^ variant had a measurable effect in cardiomyocyte response to stress. To this end, we generated two additional isogenic iPSC lines bearing the rs2234962/BAG3^C151R^ variant in either heterozygosity or homozygosity. This way we would be able to study gene-dose effects. We exposed iPS-CMs from these cell lines to increasing doses of the proteasomal inhibitor bortezomib, a well-known cardiotoxic drug^36–38^ to which BAG3-deficient cells are particularly sensitive^25, 34^. Interestingly, iPS-CMs bearing BAG3^C151R^ variants showed a higher EC_50_ than BAG3^WT^ cardiomyocytes did, indicating a greater resilience to the inhibitor. This effect was stronger in BAG3^C151R^ homozygous cardiomyocytes than in heterozygous ones, suggesting a response influenced by gene dose ***(Fig 3A, Fig S7A)***. By contrast, cardiomyocytes expressing the pathogenic BAG3^E455K-3xFLAG^ variant were significantly more susceptible to bortezomib than BAG3^WT^ cardiomyocytes ***(Fig 3A, Fig S9A)***, and had an EC_50_ similar to that of BAG3^-/-^ cell lines, consistent with observations that pathological BAG3 mutations that impact the BAG domain can have a dominant negative effect^29^. We also noted that BAG3 overexpression was able to rescue the BAG3^-/-^ phenotype and provide an additional level of resistance over BAG3^WT^ ***(Fig S7A-B)***, in agreement with studies using BAG3 overexpression as a potential cardioprotective strategy^39, 40^.

**Figure 3.**
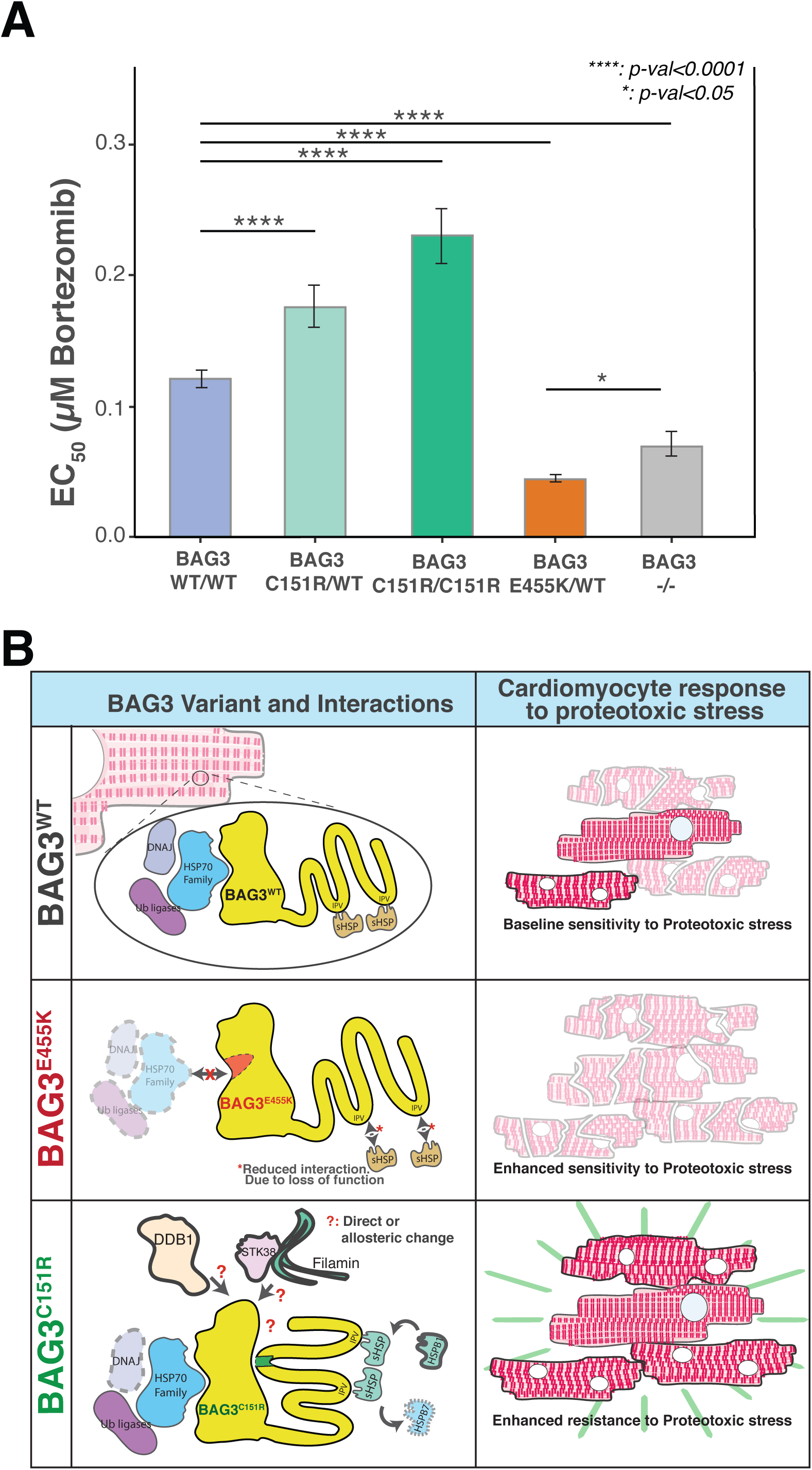
The rs2234962-BAG3^C151R^ variant provides enhanced resistance to the proteotoxic drug bortezomib in iPS-CM. (A) Calculated EC_50_ and 95% confidence intervals for Bortezomib in control (WT/ WT), BAG3^C151R^ heterozygous (C151R/WT) and BAG3^C151R^ homozygous (C151R/C151R) iPS-CMs. Cells bearing the pathogenic variant BAG3^E455K^ (along with a 3xFLAG epitope) and cells lacking BAG3 expression were used as a positive control for increased sensitivity to bortezomib. *N=*3. ****: P-value<0.0001 and *: p-value<0.5 using one-way ANOVA with post-hoc Zidak correction. (B) Graphic summary of the data presented in this study: BAG3^C151R^ changes the profile of BAG3 protein interactions (mediated by a hypothetical structure conformational change or stabilization of a particular structural state or transition) toward factors required for cardiomyocyte myofibrillar quality control (DDB1, STK38, Filamin). This in turn results in an enhanced resistance to proteotoxic insult. It is unclear if the increased binding of myofibril quality control factors is driven by a direct interaction with the local amino acid sequence or by an allosteric effect.

Finally, to rule out that the change in cardiomyocyte response to stress was due to an increase in BAG3 protein levels, we quantified protein expression in iPS-CM from the cell lines bearing BAG3^C151R^ and BAG3^E455K^ variants; we found no significant changes in protein levels, compared to BAG3^WT^ ***(Fig S9C)*.**

These results show that the BAG3^C151R^ variant provides an enhanced, dose-dependent resistance to proteotoxic stress in cardiac muscle cells *in vitro*, and that a single amino acid change rather than a change in protein levels is responsible for this effect. A proposed mechanism for the influence of single nucleotide variants on BAG3 interactions and iPS-CM response to stress is show in ***(Fig 3B)*.**

## Discussion

In this study, we report that the common variant rs2234962—which results in the BAG3^C151R^ amino acid change—causes a change in the interaction profile of the BAG3 protein and is directly associated with a heightened resistance of human cardiomyocytes to proteotoxic stress. Although numerous studies have associated the haplotype block containing this variant to a lower incidence of DCM, direct causality had been hard to infer from genetic association studies alone, in part due to the multiple linked nucleotide variants in this region, and the lack of physiological relevant models to study such cardioprotective effects. Since increasing BAG3 protein levels has been shown to have a cardioprotective effect^39, 40^, we considered two hypotheses: 1) that the C151R amino acid change directly modifies BAG3 function in a manner leading to cardioprotective effects and 2) that the haplotype block functions primary by regulating BAG3 expression. Here, we sought to test the first hypothesis. We utilized genome engineering to precisely insert the BAG3^C151R^ variant in an otherwise unchanged genetic background, and used disease modeling powered by iPSCs to identify a protective effect on cardiomyocytes exposed to a cardiotoxic drug. Although we have not tested the effect of other genetic variants in the same haplotype block, our observation that BAG3^C151R^ is sufficient for a functional change without a significant change in protein levels strongly suggests that this coding change is driving at least some of the cardioprotective effect observed in genetic association studies. Our data also indicates an allele dosage effect for this protection, in agreement with some genetic studies that suggest homozygous individuals have an even lesser risk of disease than heterozygous individuals^5, 13^.

This study demonstrates the strength genetically engineered iPSCs to dissect the effects of human genetic variants on cardiac myopathies. The ability to perform genome engineering techniques on iPSCs and the increasing breadth of functional information that can be obtained from iPS-CMs enables prospective studies in a cost- and time- efficient manner. Here, in addition to characterizing cardiomyocyte-specific protein interactions, we used an automated machine learning pipeline to objectively analyze microscopy images from different BAG3-associated gene knockdowns. This adds a layer of information to the putative interaction list by quantifying the contribution of each one of the factors to the myofibrillar structure, allowing us to prioritize targets by identifying which ones are involved in BAG3-associated maintenance of sarcomeric integrity. A similar approach has been used to study small molecule perturbations on iPS-CMs and other cell types^41, 42^. The use of iPS-CMs as a model for cardiomyopathies is not without limitations: iPS-CMs are known to lack some important features of mature myocytes and are not exposed to mechanical stress from pressure load, which is a predominant factor in the challenging of the myocardial protein quality control machinery^28, 43–45^. This last point could influence the profile of BAG3 interactions and might explain the lower representation of structural and sarcomeric proteins in our BAG3 interactome compared to other similar studies using primary heart tissue^27, 28^. Despite the highlighted differences between iPS-CMs and mature cardiomyocytes, we and others have been able to identify functional changes in genome-edited iPS-CMs that are consistent with the observations from human genetic association studies^25, 34, 44, 45^. Here, we used the proteasome inhibitor bortezomib as an inducer of proteotoxic stress. Bortezomib is a chemotherapy drug with a marked cardiotoxic effect, which acts by stalling the proteasome networks and increasing the accumulation of misfolded proteins^46^. This buildup of toxic protein species is also a hallmark of the aging and heart failure myocardium, presenting bortezomib insult as a useful tool to study cardiac protein quality control. This correlation between *in vitro* results and genetic studies suggests that the role of BAG3 in iPS-CM is at least partially representative of its function in the adult human heart muscle. In addition, we were only able to identify significant changes in co-precipitation with the BAG3^C151R^ when using a cardiomyocyte background, stressing the importance of using a disease-relevant cell model for the functional study of cardiomyopathy disease variants.

The differences in co-precipitation partners between the BAG3^C151R^ and the wild type protein point towards a molecular mechanism for the cardioprotection observed in genetic association studies. The largest increase in binding was observed for Filamin A (FLNA). The gene for FLNA is expressed at high level in embryonic cardiomyocytes, but its expression goes down in the adult striated muscle in favor of a paralog, Filamin C^47, 48^. Although we also identified FLNC in our pulldown samples, the high abundance of Filamin A in iPS-CM could be facilitating an interaction in this cellular background. The importance of FLNA in iPS-CM is also highlighted by the strong phenotype observed in our gene knockdown analyses. The BAG3^C151R^ variant also showed a significant (∼2-fold) enrichment on the hippo pathway kinase STK38/NDR1, which was recently described as an inhibitor of BAG3-triggered autophagy^49^, and ubiquitin ligases DDB1 and TRIM21, which could potentially aid in the BAG3-mediated disposal of ubiquitinated defective proteins^26^. Both HSPB1 and STK38/STK38L have been independently described to mediate Filamin phosphorylation^50, 51^. We also observe a reduction in the co-precipitation of cardiac-specific small heat shock protein HSPB7. Although this protein has been previously described to not interact directly with BAG3^52^, we identify it in our co-precipitation purification studies, suggesting it may be an indirect interaction through other interaction partners (for example, HSPB8^53^) and that the BAG3^C151R^ change may tilt the sHSP composition of BAG3 complexes in favor of other members (such as HSPB1). Residue 151 is located in a conserved region of the BAG3 protein with no known domain or function (see Figure S1A). Due to its proximity to the IPV domains, which mediate interaction with sHSPs^54^, the BAG3^C151R^ transition could be directly affecting the binding of these. Alternatively, due to the predicted intrinsically disordered nature of this segment of BAG3, it is possible that such change could have an effect on distant regions of the protein, such as the BAG domain that is responsible for indirectly binding of BAG3 protein targets for quality control.

It is particularly interesting that knockdown of the BAG3^C151R^ differential interactors scored as being closely related to BAG3 insufficiency in our myofibrillar staining image analyses. BAG3 is a known protein hub that dynamically connects different subcellular processes resulting in a broad functional spectrum^21, 54^. We propose that the BAG3^C151R^ protein variant tilts the balance of interactions to favor a number of factors that are required for the maintenance of myofibrillar structure in a BAG3 dependent manner (FLNA, DDB1, STK38, HSPB1 among others) (Fig 3B). This could in turn lead to enhanced function by either interaction stabilization or more effective detection and disposal of misfolded species. Future studies will be required to test these and other possible models for the BAG3^C151R^ mechanism of action.

To our knowledge, the data presented here provides the first evidence of a functional change associated with the rs2234962/BAG3^C151R^ variant, and the first time a putatively protective hit from a heart failure genetic association study has been validated *in vitro*. In the human heart, cardiomyocytes are under constant physical stress and have very limited proliferative potential, making them rely on an active proteostasis network to properly assemble and dispose of myofibrillar factors. The BAG3 co-chaperone is a key member of this machinery, and loss of protein expression or loss-of-function variants are known to lead to dilated cardiomyopathy and heart failure. Our study shows that a specific amino acid change in the BAG3 protein can also lead to the opposite effect, cardioprotection. Despite the high frequency of BAG3^C151R^ (∼20% in European and South-East Asian populations), no negative health effects have been described to date for this variant. This opens an avenue for further exploration of therapeutic strategies that tackle the role of BAG3 in the heart, particularly those aiming to mimic the effect of the BAG3^C151R^ variant, as a way to improve cardiac muscle function and aid in the treatment or prevention of hereditary and acquired heart failure.

## Methods

### BAG3 genetics and conservation plots

Linkage disequilibrium plots for the BAG3 putative protective allele block were generated using data downloaded from the International Genome Sample Resource (IGSR) database^55^, using the data for the CEU population (Utah residents with Northern and Western European ancestry). The map graphs with the geographic distribution of allele frequencies for the rs2234926 variant was obtained using the Geography of Genetic Variants (GGV) browser tool^56^ and edited in-house to enhance readability. Both tools used population data from the 1000 Genomes Consortium^57^. Aminoacid conservation data and zoomed-in plots were obtained from Aminode database^58^.

### iPSC culture and differentiation into iPS-CM

The WTC iPSC line was derived from a healthy male volunteer and is widely used as a normal control. WTC and derived cell lines were maintained and differentiated as previously described^25^. Briefly, for routine maintenance, cells were cultured on Growth Factor Reduced Matrigel (8 μg/ml, BD Biosciences) and maintained in mTesr1 medium (STEMCELL Technologies) every day. Whenever the cells reached 70-90% confluence, they were harvested using Accutase Cell Detachment Solution (STEMCELL Technologies) or ReLeSR Human Pluripotent Stem Cell Passaging Reagent (STEMCELL Technologies), and maintained on media supplemented with the ROCK1 inhibitor Y-27632 (10 µM, Selleckchem) for the first 24 hours. Cardiomyocyte differentiation from iPSCs was performed as previously described^25^. Briefly, iPSCs were seeded on Matrigel-coated plates, at a density between 6.25 x 10^3^ and 2.5 x10^4^ cells/cm^2^. When cells reached 40-80% confluence (day 0), media was changed to RPMI1640 (Gibco) with B27 supplement (without insulin; ThermoFisher)) containing 6 μM or 12 μM CHIR99021 (Tocris) for 24 or 48 hours. Initial seeding, confluence level at day 0 and optimal concentration and exposure time for CHIR were optimized for each line. Cells were then incubated in RPMI/B27 (-Insulin) for 24 hours before changing media to 5 μM IWP2 (Tocris) in RPMI/B27(-Insulin). After another 48 h, media was changed to RPMI/B27 containing insulin. Fresh RPMI/B27 was exchanged every 3–4 days thereafter, and cells were monitored for beating daily.

### iPS-CM enrichment and cryopreservation

At day 15, differentiation cultures that showed beating in >20% of surface area were harvested using 0.25% Trypsin (Gibco) and replated. For enrichment of iPS-CMs, we used a metabolic selection protocol as previously described ^25^. Briefly, cells were allowed to recover for 72 hours, and then media was thoroughly washed with PBS and replaced with DMEM (no glucose, with sodium pyruvate, ThermoFisher Scientific) supplemented with Glutamax, Non-Essential Amino Acids, and buffered lactate (4 mM). Cells were treated with lactate media for 48 hours, three consecutive times. Then cells were allowed to recover in RPMI/B27(+Insulin). For each differentiation batch, a separate well was prepared in parallel to be used for cell counting, quantification of differentiation efficiency and genotyping. Differentiation efficiency was quantified using flow cytometry as described previously ^59^ using mouse monoclonal antibody against cardiac troponin-T (clone 13-11, Thermo Fisher Scientific). All iPS-CM cultures used in this study were >85% cTnT+, day 30. For cryopreservation, day-30 enriched iPS-CM cultures were harvested using 0.25% Trypsin and stored in CryoStor Cell Freezing Medium (STEMCELL Technologies) in liquid nitrogen. For thawing, cells were plated in thawing media (RPMI/B27(+Insulin) supplemented with 10 μM ROCK inhibitor and 20% HyClone Fetal Bovine Serum (Fisher Scientific) and directly seeded onto cell culture plates as desired. Two hours later, media was replaced and cells were maintained as described above.

### Genome engineering of iPSC lines

Derivation of the BAG3^-/-^, BAG3^WT-FLAG^ and BAG3^C151R^ (no 3xFLAG fusion) from the WTC cell line has been described previously^25, 60^. For the generation of the BAG3^E455K^ cell line, we introduced the corresponding single nucleotide mutation in the 3xFLAG donor plasmid using QuikChange Site-Directed Mutagenesis Kit (Agilent). Then this donor plasmid was used for the same process used to generate the BAG3^WT-FLAG^ line^25^. To generate the inducible expression cell line BAG3^-/-^: TetON-BAG3^WT-3xFLAG^, we used a TALEN pair described elsewhere^61^. Two million WTC iPSC were nucleofected using an Amaxa nucleofector 2B and the Nucleofector Kit C (both from Lonza), 0.5 µg of each AAVS1 TALEN pair and 1 µg of the donor plasmid (see Figure S4). Cells were then selected on 1 mg/ml G418, before picking single clonal populations for genotyping. To generate the BAG3^C151R/WT^ and BAG3^C151R/C151R^ knock-in cell lines, 8x10^5^ WTC iPSC were nucleofected using Amaxa nucleofector 2B and Solution P3 (Lonza), 80 pmol Truecut v2 Cas9 (Life Technologies), 24 0pmol sgRNA (Synthego; ACACUGUUUAUCUGGCUGAG), and 400 pmol single-stranded donor (IDT; ctcagaggtcccagtcacctctgcggggcatgccagaaacaactcagccagataaacagcgtggacaggtggctgcagcggcggcagcccagcccccagcctcccacggacctgagg). A list of all the cell lines generated and used in this study can be found in ***Table MM.1***.

**Table MM.1.**
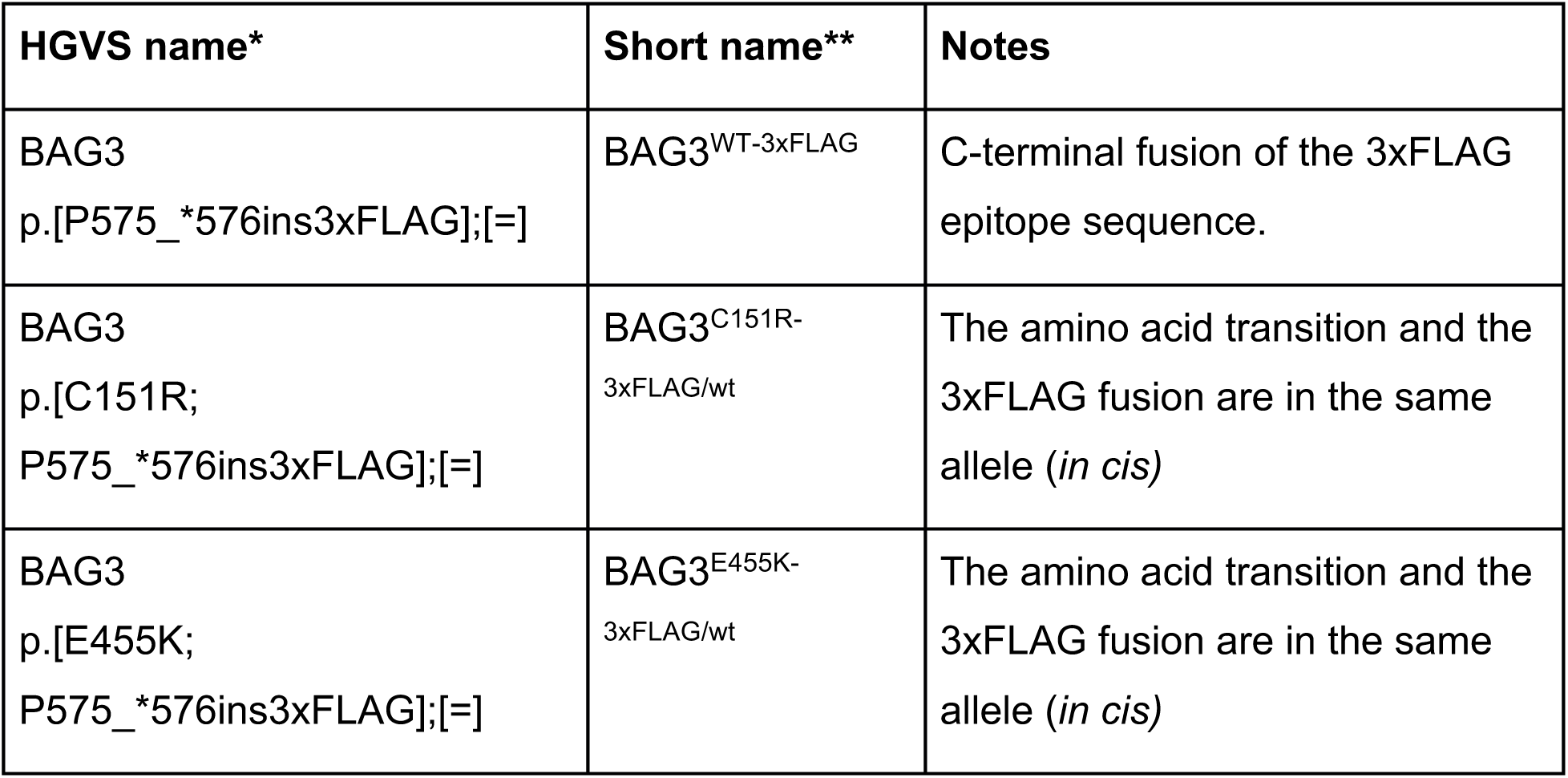
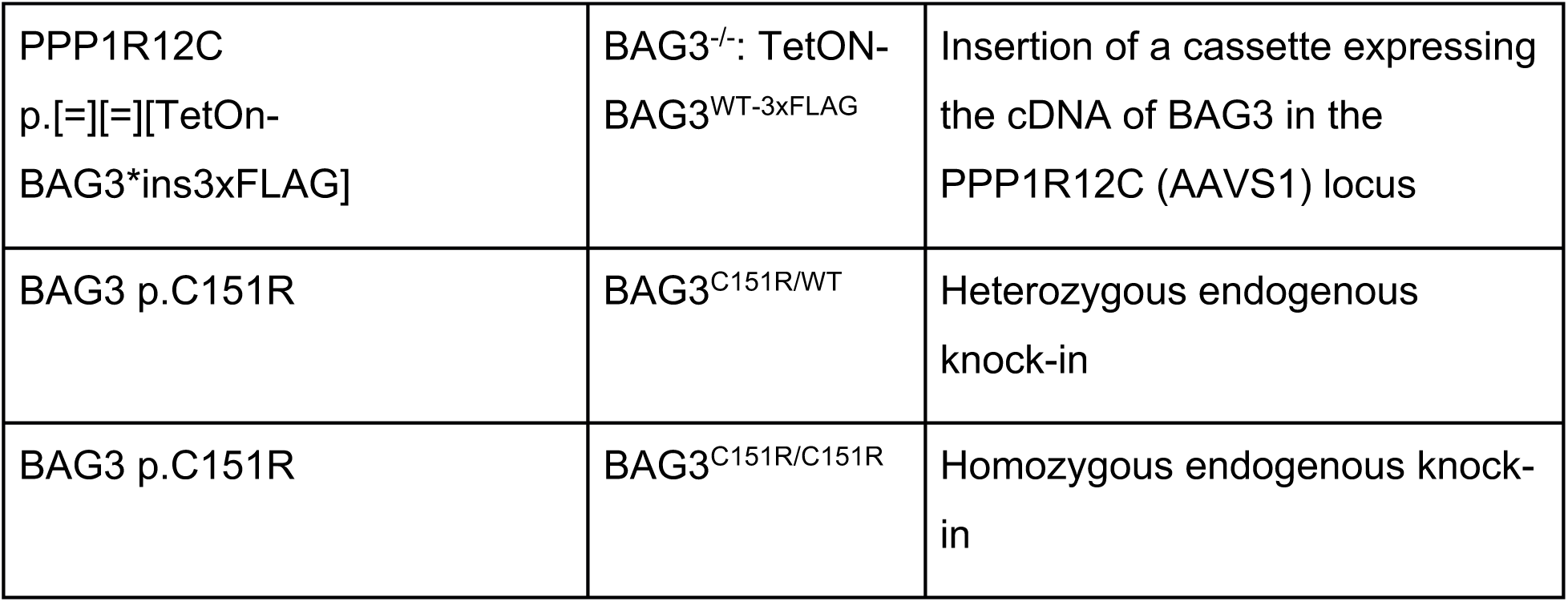
Cell lines generated in this study. *: HGVS nomenclature as per guidelines of the Human Genome Variation Society^62^. **: Short name used throughout this manuscript, based on protein level changes.

### Cell line genotyping, ddPCR linkage assay and karyotyping

For sequencing validation of genotypes, target genomic regions were amplified by PCR with BioMix Red (Bioline), sent for Sanger sequencing by MCLAB (South San Francisco, California) or Quintara Biosciences (South San Francisco, California). Quantification of the allelic abundance of the modifications inserted and verification of the excision of selection cassettes was performed by droplet digital PCR (ddPCR). For each ddPCR reaction, 50 or 100ng of genomic DNA was mixed with 5 μM of each one of two TaqMan MGB detection probe (FAM or HEX/VIC), 18 μM of forward and reverse amplification primer, and 1x of ddPCR Supermix for Probes (Bio-Rad). This mix was processed into droplets using a QX100 Droplet Generator (Bio-Rad) and the emulsion was transferred into a 96-well plate for thermal cycling. Amplifications were performed on a C1000 Thermal Cycler (Bio-Rad), using the following settings: step 1, 95°C for 10 minutes; step 2, 94°C for 30 seconds; step 3, annealing temperature (optimized for each probe pair) for 30 seconds; repeat steps 2-3 39 times; then step 4, 98°C 10 minutes. Fluorescence readouts for each droplet were obtained using a QX100 Droplet Reader (Bio-Rad) in the “absolute quantification” setting. To estimate the abundance of single nucleotide variant alleles, probe pairs were designed to discriminate between original and edited sequence. A ratio of modified vs original positive droplets was used as a readout (i.e. ∼1:1 for heterozygous). For copy number variation analysis of the 3xFLAG and puromycin cassettes, the ratio of droplets positive for these sequences versus droplets positive to a reference gene (RPP30 PrimePCR™ Probe Assay, BioRad) was used as a readout. Primers and probes for ddPCR assays were designed using the TaqMan MGB Allelic Discrimination option in Primer Express 3.0 software (Life Technologies). Sequences of the primers used in the genotyping reactions are described in ***Table MM.2***.

**Table MM.2.**
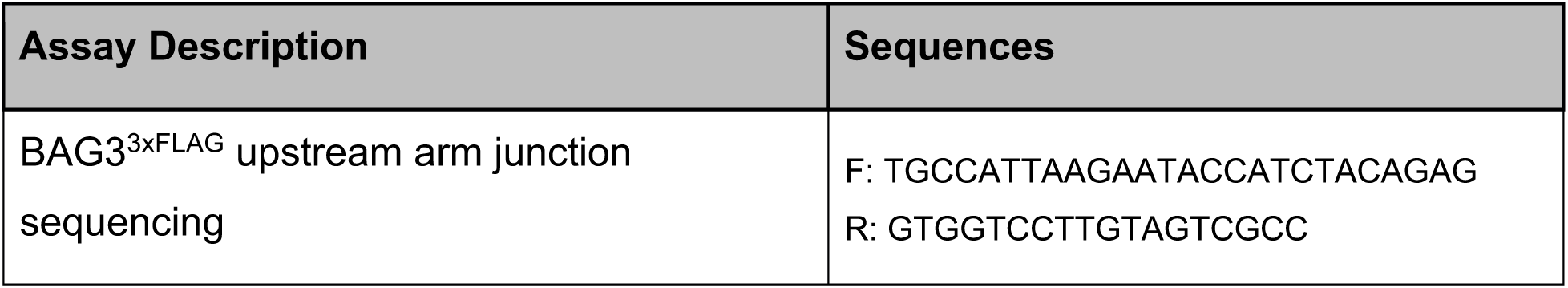
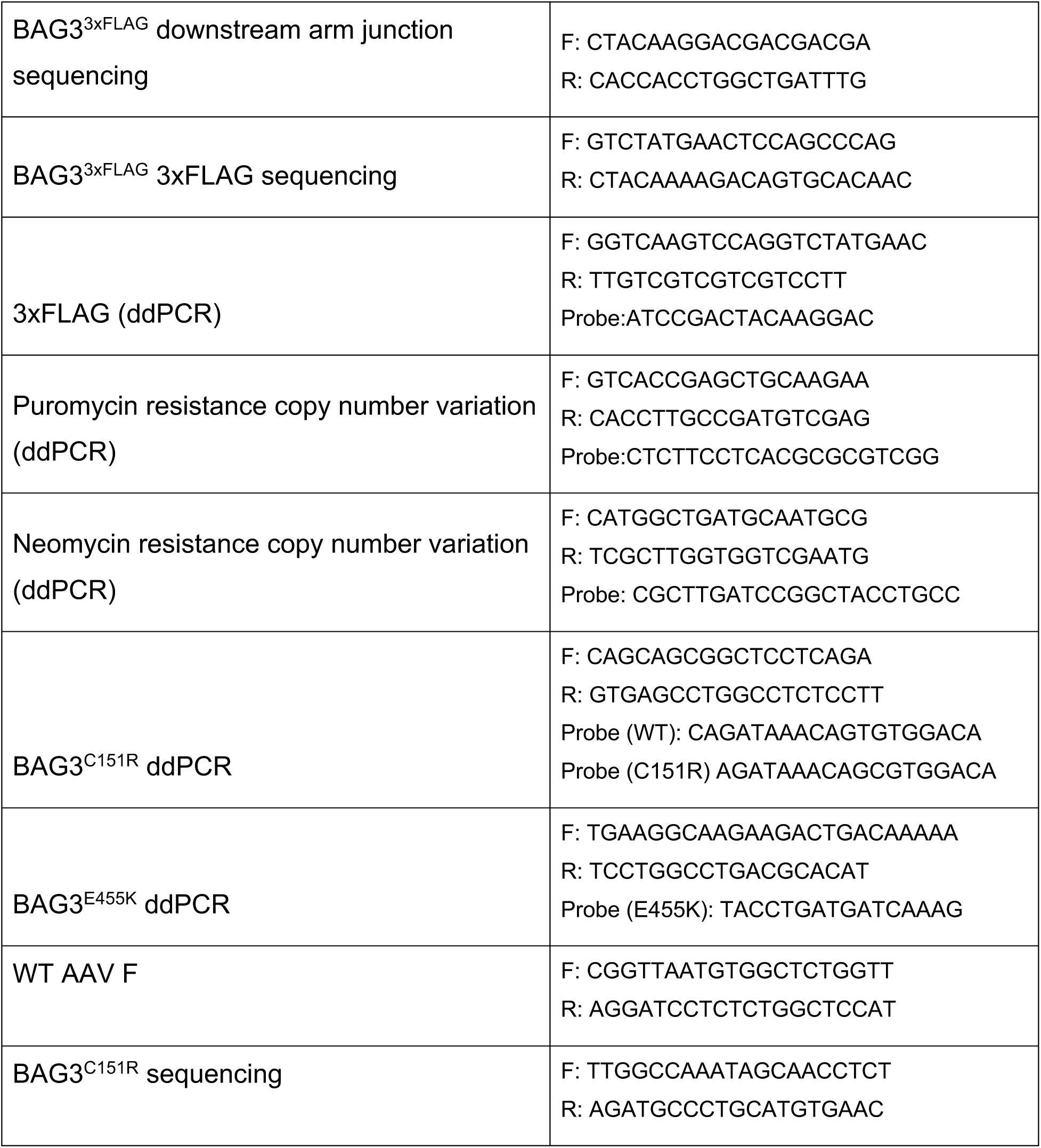
Sequences of primers and probes for genotyping reactions

For allelic phasing of the BAG3 variants and the 3xFLAG sequence, a ddPCR-based assay was used (See Fig S2). This assay combined a TaqMan probe detecting the 3xFLAG cassette and a probe binding the single-nucleotide variant we wished to phase. Droplet digital PCR was performed as described above, and the linkage percentage for variants was computed as described elsewhere ^63^.Briefly, assuming independence between events, then: 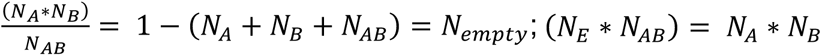, where N_A_ and N_B_ are the number of droplets positive for event A and B, respectively, while N_AB_ are double-positive droplets due to chance and N_E_ are empty droplets. If the case where the two events are physically linked is N<u>_AB_</u>, and using the Poisson statistics equation for where where 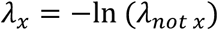, we obtain: 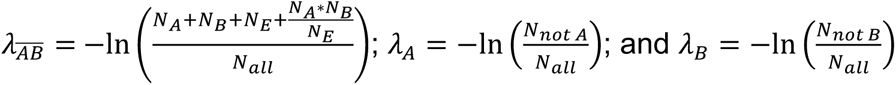. The estimated % of linked molecules is then calculated as: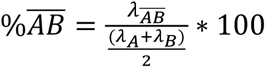.

Karyotyping of the cells was performed by Cell Line Genetics LLC (Madison, Wisconsin). Only cell lines with apparently normal karyotype were used in this study.

### Western blot and capillary immunoassay

Cells were lysed on-plate by adding RIPA buffer to the plate, incubating on ice for 30 minutes and scraping. Then lysates were harvested, clarified by centrifugation for 10 minutes and quantified using a microBCA Protein Assay Kit (Thermo Scientific). For membrane based western blot, procedure was described previously ^25^. Bands were analyzed and quantified using the ImageStudio Software (Li-Cor). All protein intensities were normalized by a housekeeping gene loading control (GAPDH). Secondary antibodies were Goat Anti-Mouse IRDye 680LT (dilution 1:20000) and Donkey Anti-Rabbit IgG IRDye (dilution 1:10000) (both by Li-Cor). For capillary immunoassay, the Simple Western Wes platform (ProteinSimple) was used following the protocol from the provider. Briefly, 1 µL of protein was diluted with Sample Buffer and mixed with Fluorescent Master Mix and denatured for 5 minutes at 95°C. For secondary antibodies, a mix of anti-rabbit and anti-mouse solutions (1:1) Secondary HRP Antibodies was used. Then all solutions were loaded on a capillary cartridge as per manufacturer’s instructions. Band intensities were quantified using Compass software (ProteinSimple) and values were normalized by GAPDH as a loading control. Primary antibodies used in this study: Anti-BAG3 rabbit polyclonal (ProteinTech, 10599-9-Ap), Monoclonal ANTI-FLAG M2 (Sigma-Aldrich, F1804), polyclonal Anti-GAPDH rabbit (Abcam, ab9458), anti-HSPB8 (Abcam, ab96837).

### HEK293T cell culture, protein overexpression and APMS

Immortalized Human Embryonic Kidney cells (HEK293T) were cultured in DMEM (high glucose, L-glutamine, sodium pyruvate; Gibco) supplemented with 10% Hyclone Fetal Bovine Serum (GE Life Sciences), GlutaMax (Gibco) and 1-00U/ml of Penicillin/Streptomycin (Gibco). Cells were passaged using 0.05% Trypsin solution (Gibco) whenever necessary. Plasmids expressing BAG3 variants were generated from BAG3 cDNA, cloned into an expression plasmid driven by CMV promoter and modified using QuikChange Site-Directed Mutagenesis Kit (Agilent). For each affinity purification, a ∼80% confluent 15 cm^2^ cell culture dish was transfected with 10 μg of plasmid using the PolyJet In Vitro DNA Transfection Reagent (SignaGen) following provider’s instructions. For controls, we used cells transfected with empty vector or a plasmid expressing the 3xFLAG peptide. Two days later, cells were harvested using Phosphate Buffer saline (PBS) with 10mM EDTA, washed twice with PBS, and resuspended in 1mL of lysis buffer (50 mM Tris-HCl pH 7.5, 150 mM NaCl, 0.5 mM EDTA, 0.5% NP-40 and 1x cOmplete protease inhibitor cocktail (Roche)). Samples were lysed by rotating at 4°C for 30 minutes, and cleared by centrifugation. For binding, the lysates were added FLAG M2 Magnetic Bead (Sigma-Aldrich; 30 μL of bead slurry per reaction) and allowed to bind 4°C for 2 hours. Beads were then collected using a magnetic stand and washed 3 times with wash buffer (50 mM Tris-HCl, pH 7.5, NaCl 150 mM, EDTA 0.5 mM, supplemented with 0.05% NP-40), with an extra wash with no NP-40. Then, beads were resuspended in 30 μL of elution buffer (1% Rapigest SF Surfactant (Waters), 5 mg/ml 3xFLAG peptide (synthesized by Elim Biopharmaceuticals Inc), in wash buffer without NP-40) and eluted at room temperature for 30 minutes with constant shaking.

Eluted protein samples were reduced with DTT (2.5 mM) at 50°C for 30 minutes, and alkylated with iodoacetamide (2.5 mM) for 40 minutes at room temperature in the dark. After that, 0.5 μg of sequencing grade trypsin (Promega) was added to the sample and incubated overnight at 37°C. Peptides were desalted and concentrated on ZipTip C18 pipette tips (Millipore) according to the manufacturer’s protocol. Digested peptides were resuspended in 0.1% Formic Acid and analyzed by LC-MS/MS on a Velos Pro ion trap mass spectrometer (Thermo Scientific) system equipped with an Easy-nLC 1000 ultra high pressure liquid chromatography and autosampler system (Thermo Scientific). Samples were injected onto a pre-column (2 cm x 100 μm I.D. packed with ReproSil Pur C18 AQ 5 μm particles) in 0.1% formic acid and then separated with a two-hour gradient from 5% to 30% ACN in 0.1% formic acid on an analytical column (10 cm x 75 μm I.D. packed with ReproSil Pur C18 AQ 3 μm particles). Each full scan was followed by 20 collision-induced dissociation MS/MS scans of the 20 most intense peaks. Dynamic exclusion was enabled for 30 seconds. Raw data files were converted into peak lists using PAVA software^64^. Spectra were then searched using Protein Prospector 5.10.1 (http://prospector.ucsf.edu/) using the SwissProt database of human proteins (April 2012). One missing cleavage was allowed. Fragment mass tolerance was set as 0.8 Da and parent mass tolerance as 1 Da. As for modifications, carbamidomethylation of cysteines was set as constant and acetylation of protein N-termini and methionine oxidation and methionine loss at N-termini were set as variable. The results from Protein Prospector were further filtered as follows: minimum Protein Score of 22.0, minimum Peptide Score of 15.0, maximum Protein E-Value of 0.01 and maximum Peptide E-Value of 0.05. All sample runs were manually curated for good values of bait count, total spectral counts and total number of unique proteins identified, leaving a total of 5 replicates per bait for downstream analysis. To filter out sample carryover across runs, we discarded each protein entry with less than half the spectral counts of the previous runs and that was present in less than 30% of all the experiments, similar to what others have described^65^. To deal with non-unique peptides, we took only the first protein with the most unique peptides and any other protein in the group that had at least one unique peptide across replicates. Spectral count values were then normalized to equate BAG3 counts (except for the truncation variants) and SAINTexpress v3.6.34 ^66^ was used with the sample compression option turned on to 4 samples (‘-R4’) for the statistical analysis of interactors. Control experiments were not compressed to minimize the impact of unremoved carryover. The scoring was performed independently for each bait condition. A database containing all GO terms with less than 20 members was input to boost scores of known interactors^66^. A final FDR cutoff of 0.1 was used. The dot plot for visualization of results was generated using the ProHits-Viz suite^67^. Cytoscape^68^ version 3.4.0 was used for the network graph generation, with the CORUM^69^ database (version 3.0) used for interconnecting the nodes. Output from SAINTexpress analyses of BAG3 variants expressed in HEK293T cells can be found in ***Table S1***.

### Immunoprecipitation of iPS/iPS-CM samples and unbiased proteomics analysis

At day 30 of differentiation, enriched iPS-CM cultures were harvested by scraping on ice-cold PBS, washed twice with ice-cold PBS and flash-frozen in dry ice with ethanol. For the bortezomib treatment condition, bortezomib was added to the media to 100 μM final concentration 24 hours prior to the harvesting. For iPS-CM APMS, 25-30 million cells per sample were used. For iPSC APMS, three ∼90% confluent 15cm dishes were used per sample. As a negative control for nonspecific binding, WTC iPSC or iPS-CM cells not expressing any 3xFLAG affinity epitope were used. All the experimental conditions were performed in four replicates, each one on a different day. For each replicate all the different conditions were run in parallel. For mass spectrometry, all samples were run sequentially. Cell pellets were thoroughly resuspended in lysis buffer (0.1% NP-40, 300 mM NaCl, 20% Glycerol, 2 mM MgCl2, 0.5 mM EDTA, 0.5 mM EGTA, 1 mM PMSF, 1 mM DDT in 50 mM HEPES-NaOH pH 8.0, supplemented with complete protease inhibitor cocktail (Roche) and Benzonase Nuclease (50 U/ml, Sigma-Aldrich)). Cells were lysed by four flash freeze-thaw cycles, followed by incubation for 20 min at 4°C in constant rotation. After clearing lysates by centrifugation, protein extracts were diluted three-fold (in 0.5 mM EDTA, 0.5 mM EGTA, 1 mM PMSF in 50mM HEPES-NaOH pH 8.0) to reduce salt content and incubated with 30 μL anti-FLAG M2 Magnetic Beads slurry (Sigma-Aldrich) for 2-3 hours. Beads were then rinsed in wash buffer (3x washes in 0.01% NP-40, 1 mM PMSF, 0.5% EDTA, 0.5% EGTA in 50 mM HEPES, pH 8.0; 1x wash on buffer without NP-40). Beads loaded with FLAG-enriched proteins were reduced (5 mM TCEP), alkylated (15 mM iodoacetamide), and digested with 1% (w/v) trypsin overnight. Resulting peptides were desalted by OMIX C18 desalting tips (Agilent) following protocol by provider and dried on a speed-vac. Peptides were resuspended in 0.1% formic acid and an iRT peptide standard mix (Biognosys) was spiked in prior to use for mass spectrometry.

For unbiased (non-targeted) analysis of pulldown samples, peptides were analyzed by liquid chromatography tandem mass spectrometry (LC MS/MS) with an Easy-nLC 1000 (Thermo Fisher, San Jose, CA) coupled to an Orbitrap Fusion Tribrid Mass Spectrometer (Thermo Fisher Scientific, San Jose, CA). Online LC separation was carried out using a 75 µm x 25 cm fused silica IntregraFrit capillary column (New Objective, Woburn, MA) packed in-house with 1.9 µm Reprosil-Pur C18 AQ reverse-phase resin (Dr. Maisch-GmbH). Peptides were eluted at a flowrate of 300 nL/min using a linear gradient of 5–30% B in 45 min, and 30–95% B for 25 min (mobile phase buffer A: 0.1% formic acid; mobile phase buffer B: 0.1% formic acid in ACN). Survey scans of peptide precursors from 400 to 1600 m/z were performed at 120K resolution in the Orbitrap, with an AGC target of 2×105, and a maximum injection time of 100 ms. Tandem MS (MS2) was performed by isolation with the quadrupole, HCD fragmentation with normalized collision energy of 30%, and rapid scan MS analysis in the ion trap. The MS2 ion count target was set to 10^4^ and the max injection time was 35 ms. Precursors with charge state 2–7 were sampled for MS2 and dynamically excluded for 20 s (tolerance of 10 ppm). Monoisotopic precursor selection was turned on, and the instrument was run in top speed mode with 3-s cycles. For protein identification and quantification, MaxQuant software v1.5.3.30 was used^70^. Tandem mass spectrometry (MS/MS) spectra were searched against the November 2016 release of the UniProt complete human proteome sequence database. MaxQuant was run on default parameters, allowing for 2 maximum missed cleavages, with a first search peptide tolerance of 20 ppm and a main search peptide tolerance of 4.5 ppm. Methionine oxidation and N-terminal acetylation were set as variable modifications, and carbamidomethylation of cysteines as fixed modification. The ‘match between runs’ setting was activated (window of 0.7 min) to improve peptide identification. Mass spectrometry raw data and search results files have been deposited to the ProteomeXchange Consortium via the PRIDE partner repository^71^.

Peptide sequences were used to validate pulldown of intended BAG3 variants. Proteins with two or fewer peptides identified were discarded, as were typical common contaminant proteins (downloaded from http://maxquant.org/). Intensity values were normalized by median equalization of the peptide lists, and aggregated into protein intensity values. SAINTq v0.0.4^72^ was then used to compare each experimental condition to the no FLAG control set individually. Significant putative protein interactors were selected with a Bayesian False Discovery Rate (FDR) cutoff of 0.1. Euler diagrams were generated using BioVenn^73^ or the R package *eulerr* ^74^.

### Targeted proteomics follow-up of iPS-CM APMS high confidence protein-protein interactions

A list of 118 proteins from our unbiased APMS study was chosen for targeted proteomics quantitation (43 proteins significant in our iPS-CM APMS studies, 37 significant in iPSC or HEK293T cell APMS, 35 nonsignificant but previously described as interacting with or related to BAG3, 3 low-variability controls, plus BAG3 and the iRT peptides). To increase the number of peptides used for quantitation of these proteins, we downloaded extra peptide sequences and iRT standardized retention time values from SRMatlas ^75^ (www.mrmatlas.com). The isolation list of all peptides (empirical + predicted) was then used to perform a preliminary scan of a pooled sample from all conditions using a Q-Exactive Plus Mass Spectrometer (Thermo Fisher). Skyline v4.2^76^ was used to manually select best peptides and fragments based on ppm, q-value and uniqueness to protein target. The final isolation list contained 4 peptides per protein (except for BAG3, which kept 8 peptides, and the 11 iRT peptides; ***Table S3***) and was split into two acquisition methods to minimize the number of concurrent peptides per window. For targeted quantitation of peptide abundance, we used a Parallel Reaction Monitoring^77^ method on a Thermo Fisher Q Exactive. Online LC separation was carried out using a 75 µm x 25 cm fused silica PicoTip capillary column (New Objective, Woburn, MA) packed in-house with 1.9 µm Reprosil-Pur C18 AQ reverse-phase resin (Dr. Maisch-GmbH). Peptides were eluted at a flowrate of 300 nL/min using a linear gradient of 7–36% B in 52 min, and 36–85% B for 7 min (mobile phase buffer A: 0.1% formic acid; mobile phase buffer B: 0.1% formic acid in 80% ACN). Peptides were directly injected into the mass spectrometer in positive ion mode over the course of a 75 min total acquisition time. MS2 scan were acquired in profile mode for peptides defined on the inclusion list at a resolution of 17.5K, with a 2x10^5^ AGC target, 30ms maximum injection time, a loop count of 40, an MSX count of 1, a 2 m/z isolation window, and an NCE of 27.

Raw data was loaded into Skyline to manually select best peptides and boundaries, ending on a total of 113 proteins with at least 2 good peptides. MSStats^78^ was then used to format the data and obtain a matrix of log intensities per sample. Missing data (0.7% of total data) was imputed by averaging remaining replicates for the same protein and condition. For analysis of the data, we used a hypervariate linear model analysis followed by empirical bayes testing to obtain a robust estimation of differential protein abundance across samples, while controlling for replicate batch variability. For this we used the *limma* package in R^79, 80^. First, to obtain a list of high-confidence interactions for each condition, we fit a model for condition vs no-FLAG control. Then, a second model was fit to compare the intensities for all significant hits in the BAG3^C151R^, BAG3^E455K^ or BAG3 (+bortezomib) conditions against BAG3^WT^. Adjusted p-value threshold was set at 0.05 in both steps. Mass spectrometry raw data and Skyline files have been deposited to the ProteomeXchange Consortium using the Panorama Public partner repository^81^ (link: https://panoramaweb.org/28ZWVY.url).

### Immunofluorescence staining of iPS-CM and imaging

For immunostaining of iPS-CM in culture, cells were washed with PBS and fixed with 4% Paraformaldehyde at room temperature for 20 minutes. Then cells were washed three times with PBS supplemented with 0.1% Triton X-100 (PBS-Triton) and incubated with 5% Bovine Albumin Serum in PBS-Triton for 1 hour at room temperature to block and permeabilize. Then cells were incubated overnight at 4°C with a solution of primary antibody in PBS-Triton. The morning after, cells were washed three times with PBS-Triton and then incubated in secondary antibodies in PBS-Triton for 1 hour at room temperature. Cells were washed three more times with PBS-Triton (first wash supplemented with 1:1000 DAPI) and kept in PBS supplemented with 0.02% Sodium Azide (Sigma-Aldrich) until imaging. Secondary antibodies used were Goat Anti-Rabbit Alexa Fluor 594 and Goat Anti-Mouse IgG Alexa Fluor 488 (both at a 1:500 dilution; both from Thermo Fisher Scientific). For primary antibodies, we used rabbit polyclonal anti-BAG3 (Protein Tech, 10599-1-A), mouse monoclonal anti-FLAG M2 (Sigma-Aldrich, F1804), and mouse monoclonal anti-MYBPC3 (Santa Cruz Biotechnology, sc-137180), all at a 1:200 dilution. Images used for the unbiased analysis of sarcomere structure were acquired using an ArrayScan VTI HCS Reader (ThermoFisher) at a 10x magnification. Images used for high-magnification figures in this manuscript were acquired using an ImageXpress Micro Confocal High-Content Imaging System (Molecular Devices). Raw images were processed using the R package *EBImage*^82^ or ImageJ^83^ for visualization and figure preparation.

### Gene knockdown on iPS-CM and unbiased image analysis

For quality control and protocol optimization experiments, FlexiTube siRNA (Qiagen) oligonucleotides against BAG3 or a scramble control were used. For imaging screening and validation experiments, a Silencer Select siRNA Custom Library (Ambion) was used. Gene targets were selected for being BAG3 iPS-CM interactor in our AP-MS experiments, being a prominent BAG3 interactor in the literature, or being an important myofibrillar/z-disk component or chaperone complex. For all gene targets, three different siRNA oligos were combined. High purity iPS-CM cultures (iCell Cardiomyocytes^2^, Lot#CMC331743, >99% cTnT+) were obtained from Cellular Dynamics International (Madison, WI). Vials were thawed as per manufacturer instructions and seeded on 24 well plates (for quality control analyses) or Falcon 96-well Black/Clear Imaging Microplate (Corning) at a density of ∼53,000 cells/cm^2^. Cells were allowed to recover for five days and transfected with 1 μM total siRNA using Lipofectamine RNAiMAX Transfection Reagent (ThermoFisher) (0.05 μL or 0.25 μL/well, respectively) following protocol from provider. Cells were left in the siRNA solution for 48 hours, and then media was changed for fresh cardiomyocyte maintenance media as usual. Cells were used for subsequent analyses (harvesting or immunostaining) 10 days after transfection.

For the unbiased analysis of sarcomere staining images on iPS-CM gene knockdowns, three biological replicates were performed for each siRNA condition. Screening plates included internal Scramble and BAG3 control conditions. In parallel, additional wells were prepared for each one of the BAG3 and Scramble siRNA conditions for training (32 wells) and validation (8 wells) of the model. Nine images per well were used in all conditions. Images were used to train a supervised machine learning image classification model using PhenoLearn framework (www.phenolearn.com), as described elsewhere^41, 42^. This model was used to generate a classification score for each image from the screening plates indicating the similarity to the BAG3 siRNA (score closer to 0) or Scramble siRNA (score closer to 1) training sets. For the MYBPC3 staining images, this classification score was dubbed “BAG3 Sarcomere Score”. The median score for each well was used for further analyses. Images from corner wells were discarded due to loss of viability, and conditions with less than 3 replicates were discarded. The R package *EBImage*^82^ was used for the quantification of cell viability (nuclei count) and BAG3 staining intensities. Statistical significance for the differences across conditions was performed using a One-Way ANOVA with post-hoc multiple comparisons corrected by Dunnett test. Information on gene targets selected, siRNA sequences used and image analysis summary values can be found in ***Table S2***.

### Bortezomib dose-response curve

Day 30 frozen iPS-CM vials for all the cell lines used were thawed (as described above), counted and seeded directly into 96-well plates. All lines were seeded in parallel at three different densities (15,000, 30,000, and 60,000 cells/well), each in triplicate. Cells were allowed to recover for 5 days with RPMI/B27+ media changes every other day. On day 5, media was supplemented with the desired concentration of bortezomib (Sigma Aldrich) in DMSO, with the equivalent volume of DMSO for the vehicle-only controls. Cells were incubated for 48 hours and then wells were washed twice with PBS before adding fresh RPMI/B27+ media. Seven days later, cells were treated with RPMI/B27+ with 10% resazurin blue reagent (PrestoBlue, Thermo Scientific) for 1 hour at 37°C. Fluorescence was then read for each well using a SpectraMax i3 plate reader (Molecular Devices) according to the provider’s instructions. For each cell line, the seeding density that resulted in the most comparable viability readout across lines for the DMSO-only condition was chosen for the calculations. Viability readout values were normalized to the range from 0 (media only) to 1 (vehicle only) and fit on a two-parameter log-logistic dose-response model using the R package *drc*^84^. Confidence intervals were adjusted to control for family-wise error rates using the package *multcomp*^85^. Pairwise EC50 comparisons were then performed using a One-way ANOVA test with post hoc Zidak correction using Graphpad Prism (version 8.4.3.686 for Windows, GraphPad Software, San Diego, California USA, www.graphpad.com).

## Acknowledgements

We thank the Gladstone Stem Cell Core and the Gladstone Assay Development and Drug Discovery Core for providing their technical support and experimental expertise. We also would like to thank Dr. Reuben Thomas from the Gladstone Bioinformatics core for his advice on data analysis. We are also very grateful to Dr. Jeff Johnson, Dr. Ruth Huttenhain and Dr. Gwendolyn Jang for their advice on affinity purification and mass spectrometry experiments. We thank Prof. Jason Gestwicki and their team for their scientific and technical advice. We thank Amanda Chan, Alisha Birk, Edward Shin, Chiara Marley and Serah Kang for their technical support. We also thank Francoise Chanut from Gladstone Editorial Services for her feedback on manuscript preparation.

## Author Contributions

J.A.P.B., L.M.J., P.S., N.J.K., and B.R.C. designed and supervised the study. J.P.B., C.L.J., A.T., J.J.H., W.V.R. and K.W. performed cell line generation, cell culture and differentiation. J.P.B., R.M.K, E.P. and D.L.S. performed affinity purification-mass spectrometry experiments and analyses. J.P.B., K.W. and M.A.M. performed siRNA knockdown panel and image analysis. J.P.B. and L.M.J. performed bortezomib toxicity assay and analyses. All authors contributed to writing the manuscript and preparing the figures.

---

**Figure S1.**
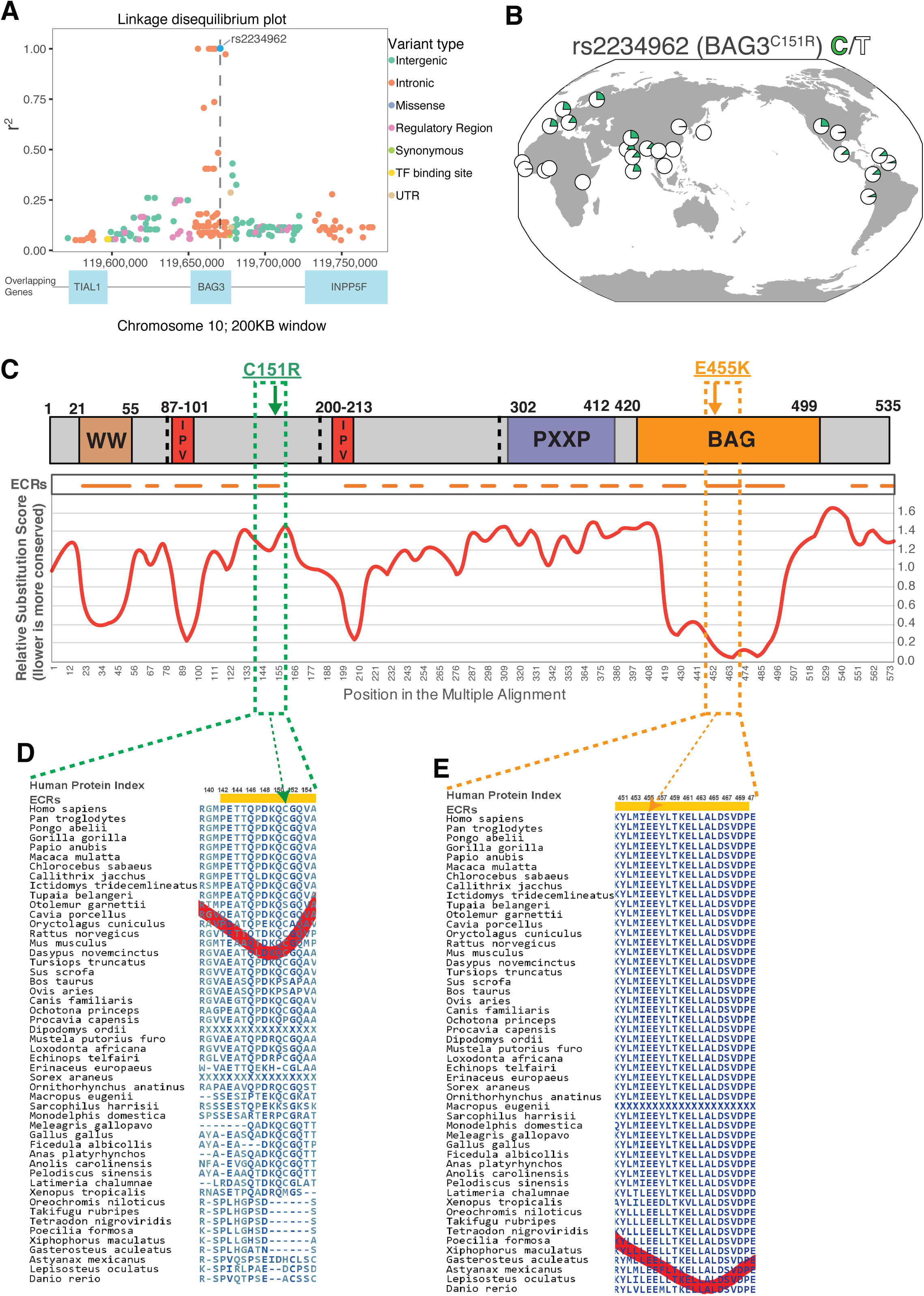
Expanded genetic data graphs for rs2234962, BAG3^C151R^. Conservation of residues affected by BAG3^C151R^ and BAG3^E455K^ variants. (A) Zoomed out version of Figure 1A, showing a window of 200KB. Dot color indicates type of nucleotide change. **(B)** Allele frequency map for rs2234962 depicting all 1000 Genomes populations. **(C)** Top: Diagram of BAG3 domain structure. Bottom: Amino acid conservation plot for matching BAG3 regions. Decreasing Relative Substitution Score regions (valleys) indicate sequences with high conservation across species and are annotated as Evolutionary Constrained Regions (ECRs). (D-E) Zoomed in regions for the ECRs around BAG3^C151^(D) and BAG3^E455^(E).

**Figure S2.**
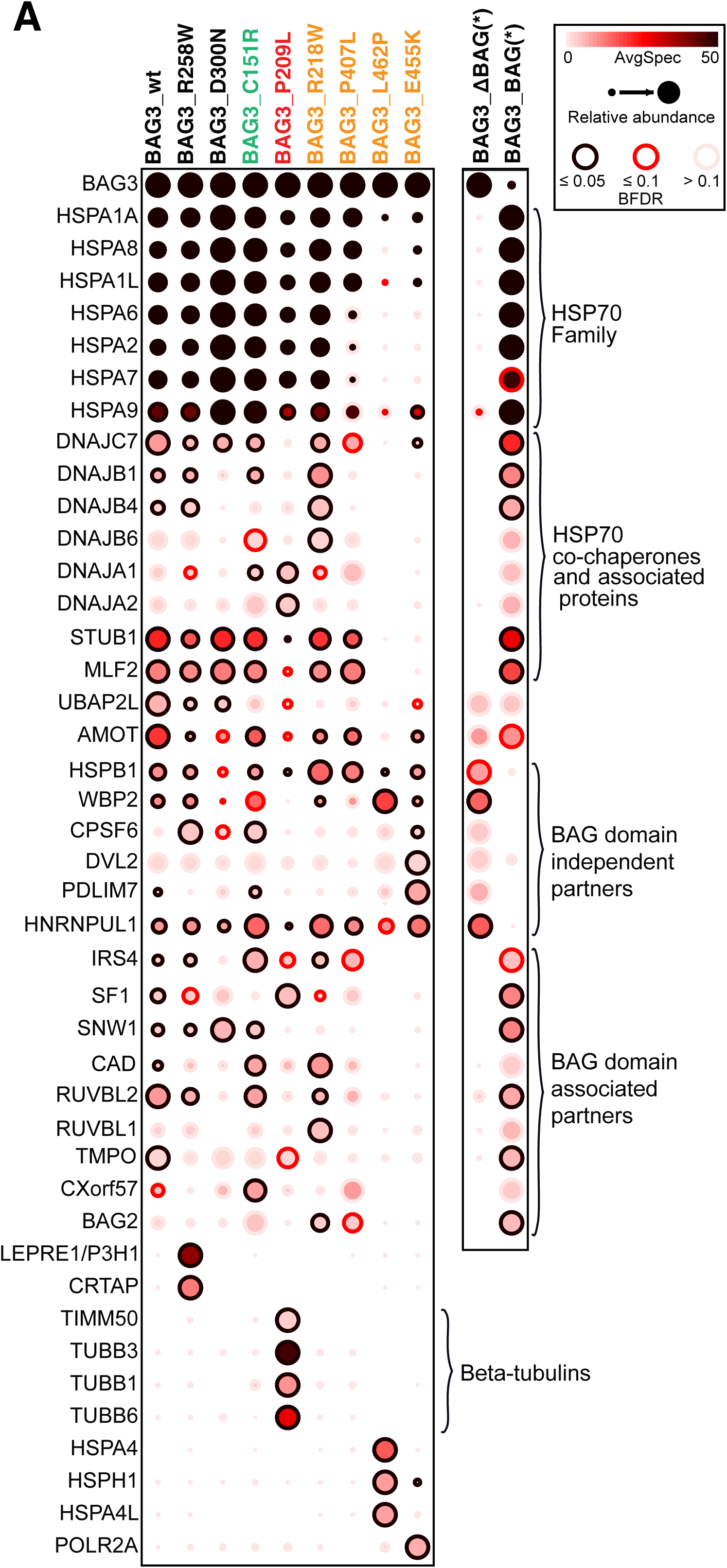
Co-precipitation profiles of different BAG3 variants overexpressed in a HEK293 cell background. Dot size represents the amount of co-precipitated protein normalized across variants. Dot color represents absolute protein abundance (spectral counts). Dot rim represents statistical significance. Yellow colored variants are known pathogenic variants associated with DCM. Green variant is putative cardioprotective variant BAG3^C151R^. Red variant (BAG3^P209L^) is associated with skeletal myofibrillar myopathy. Black variants are not associated to DCM or any other pathology. Rightmost two columns depict data for BAG3 truncated variant without the BAG domain (BAG3_ ΔBAG) and for the BAG3 protein BAG domain only (BAG3_ΔBAG). For the truncated variants, no normalization by bait levels was performed. *N*=5, BFDR obtained using SAINTexpress (see methods).

**Figure S3.**
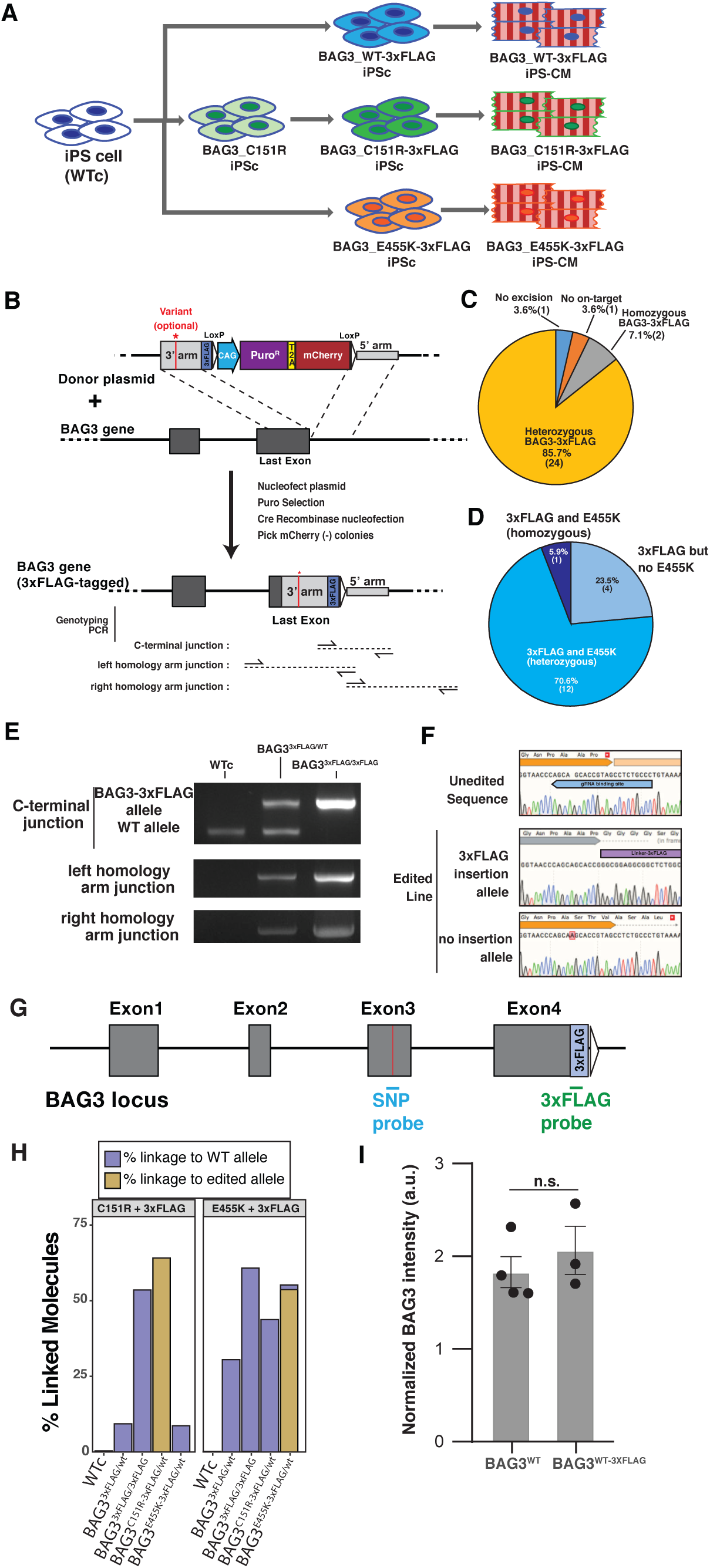
Generation of the isogenic cell lines carrying BAG3 variants and a 3xFLAG epitope tag fusion in the endogenous copy of the BAG3 gene. (A) Workflow for the cell line generation. (B) Strategy for the insertion of a 3xFLAG epitope fusion at the C-terminal of the BAG3 gene. The BAG3^C151R-FLAG^ variant was generated using the same process on a preexisting cell line bearing the C151R mutation. To generate the BAG3^E455K-FLAG^ cell line, the homology arms were engineered to contain the SNP and insert it during recombination. (C-D) Genotypes of the single-cell clones picked for 3xFLAG insertion (C) and the co-segregation of the BAG3^E455K^ variant (D). (E) Genotyping the products of the 3xFLAG insertion by PCR. (F) Cells with a heterozygous insertion of the 3xFLAG epitope tag also had a SNP in the other allele that extended the BAG3 protein product by 4 amino acids. (G-H) A droplet digital PCR phasing test was used to select clones that contained the desired SNP variants and the 3xFLAG C-terminal sequence in the same allele. The test used different probes (G) to generate an estimate of linked molecules for each cell line and probe combination (H; See Methods for more details) (I) Insertion of the 3xFLAG fusion in the BAG3 gene did not alter the protein levels. *N*=3; one-way ANOVA.

**Figure S4.**
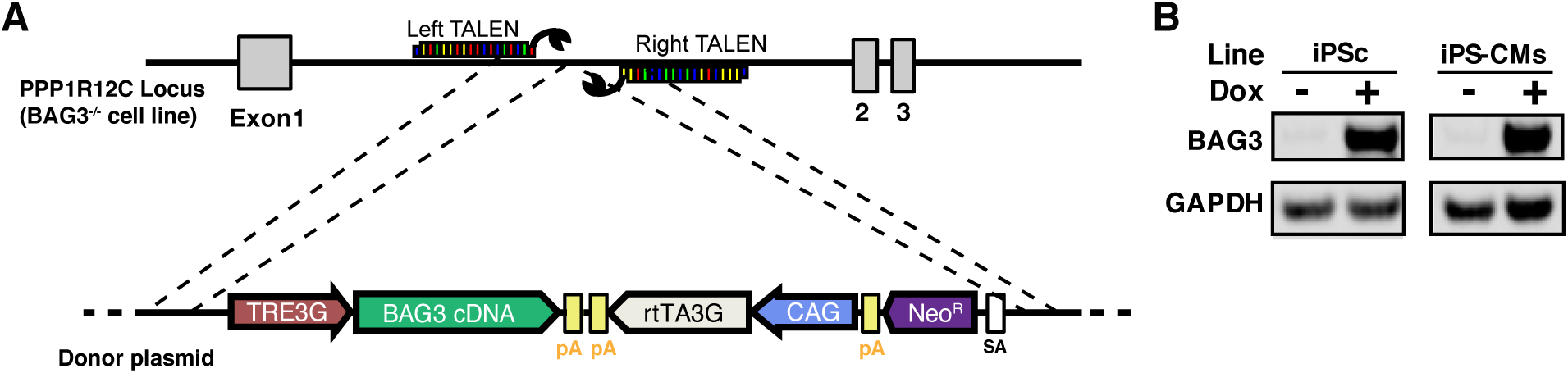
Generation of a cell line with inducible expression of the BAG3^WT^ protein. (A) Diagram of the editing strategy. On a BAG3^-/-^ cell background, a doxycycline-activated BAG3- 3xFLAG expression cassette was inserted in the PPP1R12C (AAVS1) safe-harbor locus. (B) Western blot of the BAG3 expression on BAG3^-/-^:TetOn-BAG3^WT-3xFLAG^ iPSCs and iPS-CMs with and without Doxycycline addition.

**Figure S5.**
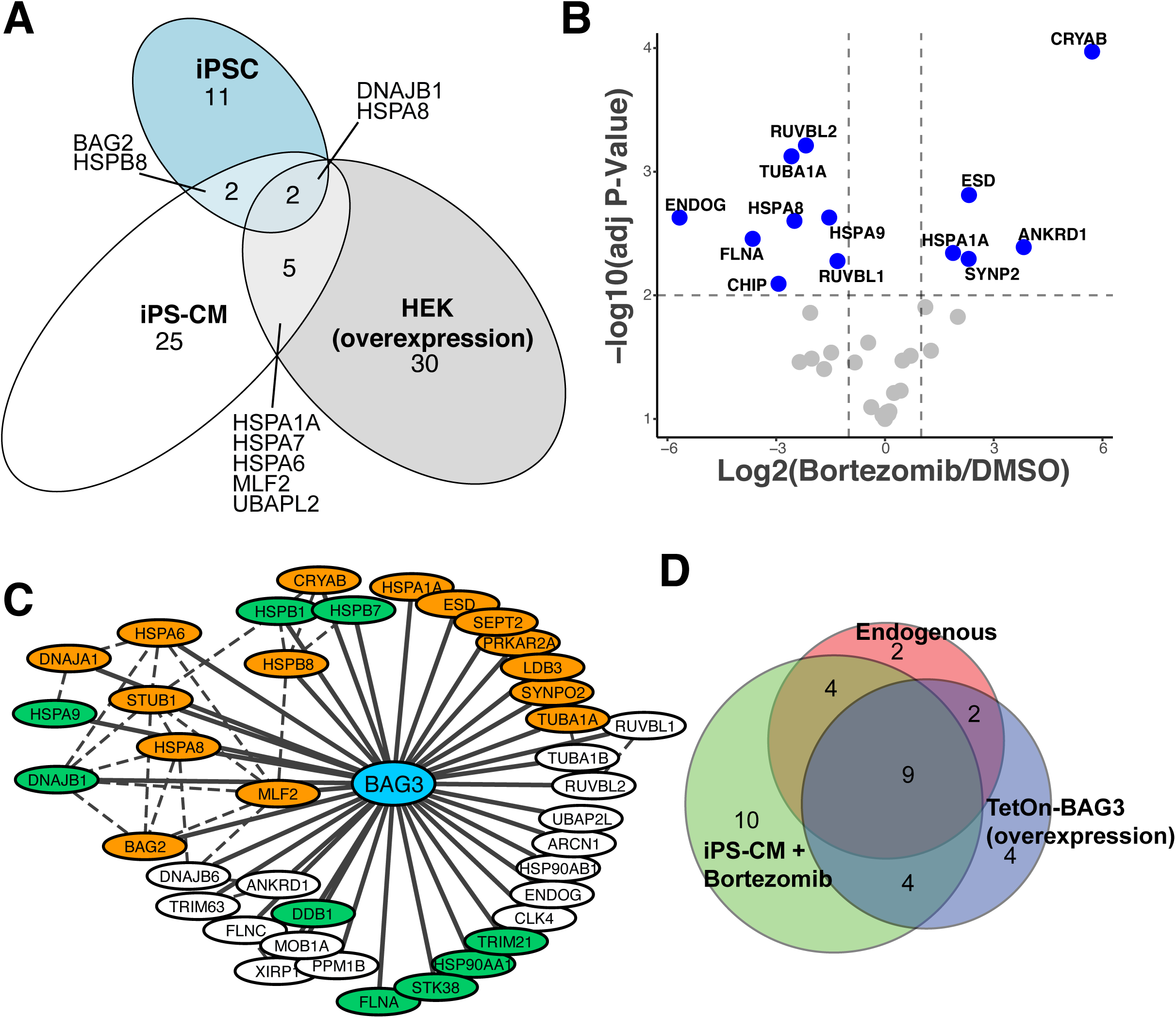
Affinity purification - mass spectrometry characterization of BAG3 binding partners in a cardiomyocyte background. (A) Venn diagram of the high confidence BAG3^WT^ protein-protein interactions identified in three cellular backgrounds. HEK293T cells had overexpressed baits, while iPSC have much lower levels of endogenous BAG3 expression than iPS-CM, which could have influenced the results. Each cell type specific dataset was scored separately against its own matched control(s). (B) Volcano plots depicting co-precipitation intensity in BAG3^WT^ cardiomyocytes treated with Bortezomib (100nM) relative to DMSO (1:10.000) for 24hours. Horizontal dashed line indicates statistical significance threshold (adjusted p-value <0.01) and vertical dashed lines indicate a fold change of 2. *N*=4. (C) Network diagram of the iPS-CM co-precipitation partners identified for BAG3 in this study. Nodes in orange indicate partners that significantly changed when pulling down BAG3^E455K^. Nodes in Green indicate partners that significantly changed when pulling down BAG3^C151R^. Dashed lines: known interactions in the iRefIndex database. (D) Venn diagram comparing the BAG3 binding partners identified in an iPS-CM background when using endogenous basal levels of expression, an overexpression system, or endogenous expression under proteotoxic stress. The graph highlights the importance of using endogenous expression for accurate characterization of binding partners, and the information gained from using a stress state.

**Figure S6.**
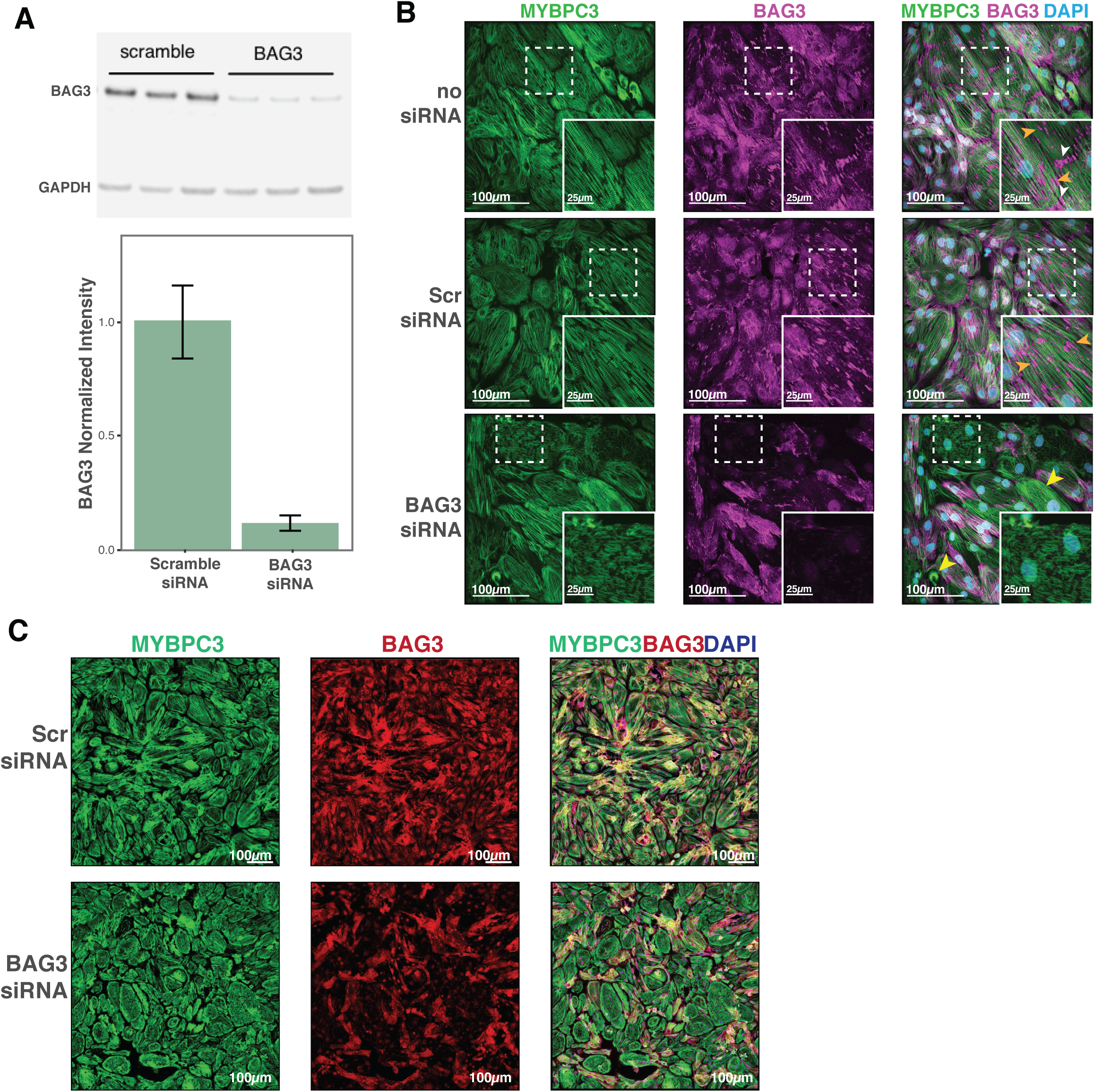
Additional sample micrographs from BAG3 knockdown iPS-CM. (A) BAG3 silencing by siRNA was effective at reducing protein levels (∼85% reduction). *N*=3. (B) Additional sample micrographs from BAG3- and Scr-siRNA-treated iPS-CM, plus a no-siRNA condition. Orange arrowheads: BAG3 accumulation on myofibrillar breaks; white arrowheads: BAG3 accumulation on polar ends of cells; yellow arrowheads: iPS-CM displaying myofibrillar aggregation and collapse. (C) Sample images in the same magnification used for the automated scoring analysis. Lower magnification allowed for faster acquisition and richer features to use directly in the scoring scheme, but higher magnification images were used elsewhere in this manuscript for easier viewing.

**Figure S7.**
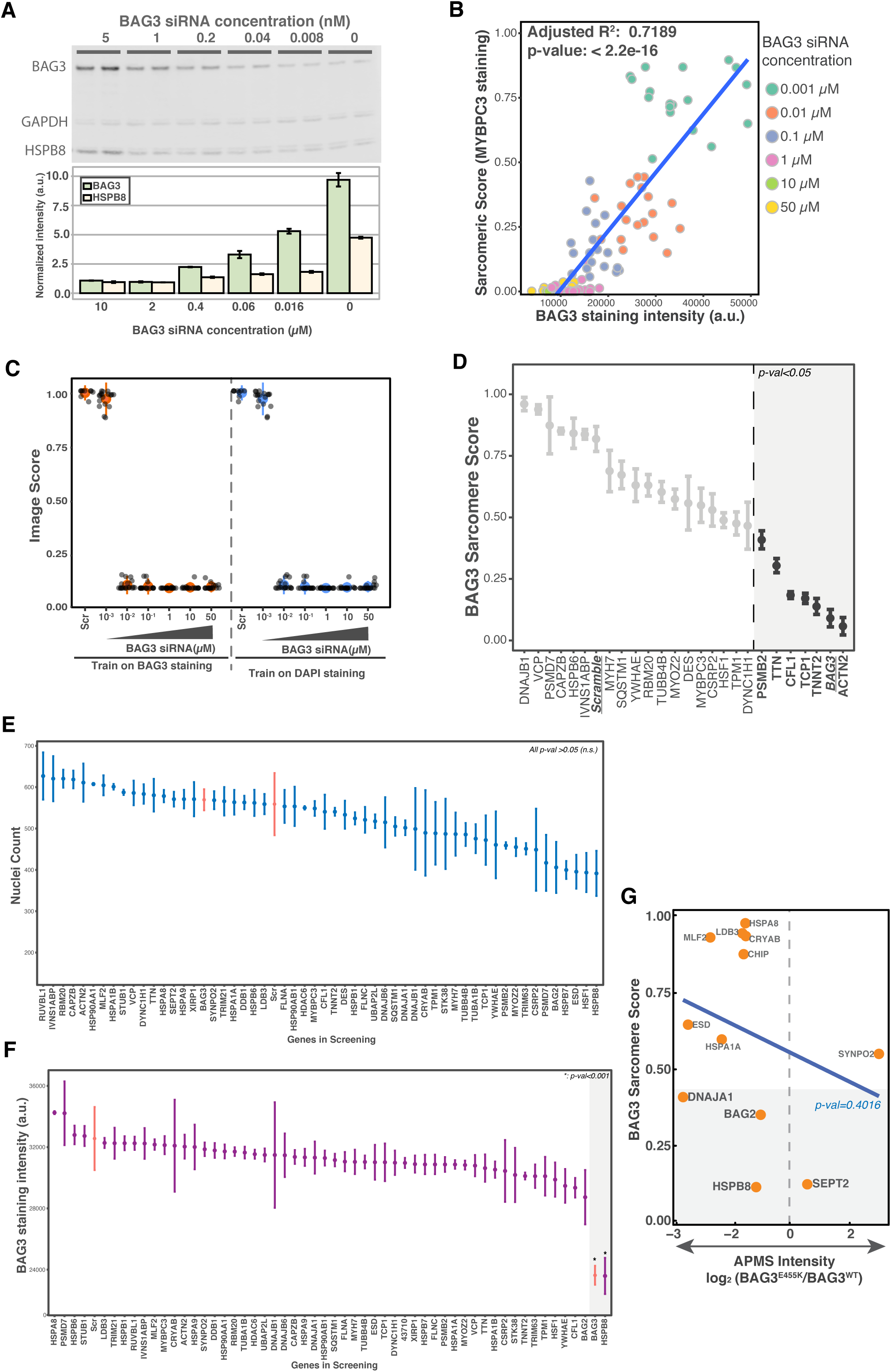
Quality control and additional data from the siRNA knockdown-myofibrillar scoring workflow. (A) Western blot demonstrating the titration of BAG3 siRNA results in decreasing cellular BAG3 protein levels. Shown at the bottom of the western blot are levels of HSPB8, which are also affected by BAG3 knockdown. *N=2.* (B) BAG3 sarcomere score inversely correlates with BAG3 protein levels. *N* = 18 images. (C) Plot of the image scores that result from training based on BAG3 or DAPI staining. These stains do not display the same dynamic range as scoring based on myofibrillar (MYBPC3) staining. *N*=9 for scramble; 18 for the rest. (D) BAG3 Sarcomere Score for the knockdown of selected factors that were not identified in our AP-MS coprecipitation studies. Dots represent mean of 3 replicates from separate wells, each being the median score of 9 images from the same well. Error bars: SEM. P-val cutoff: 0.05 using a one-way ANOVA with post-hoc Dunnett test. (E-F) Plot of the nuclei count(E) and BAG3 staining(F) intensities for the gene knockdowns used in the siRNA- myofibrillar scoring analyses. Dots represent mean of 3 replicates from separate wells, each being the median score of 9 images from the same well. Error bars: SEM. P-val cutoff: 0.05 using a one-way ANOVA with post-hoc Dunnett test. (G) Plotting of the APMS intensity ratio for BAG3^E455K^ differential interactors and their BAG3 Sarcomere Score. There is no statistically significant correlation. P-value obtained fitting a linear model. Pearson’s product-moment correlation: -0.27.

**Figure S8.**
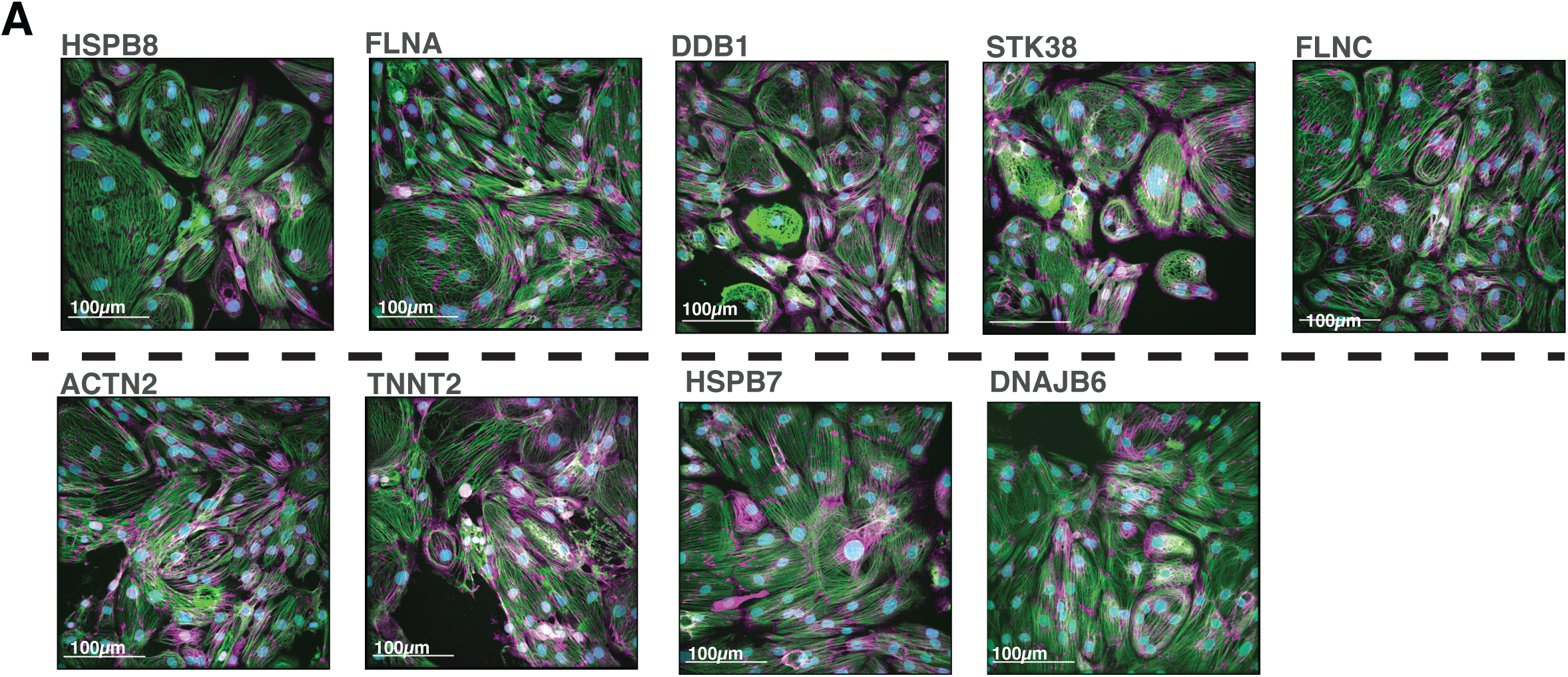
Sample images from selected siRNA knockdowns. HSPB8 knockdown was the only knockdown to significantly reduce BAG3 levels. FLNA, DDB1 and STK38 are BAG3^C151R^ differential interactors whose knockdown resulted in sarcomere scores similar to BAG3 knockdown. ACTN2 and TNNT2 are well known sarcomere components that display low sarcomere scores similar to BAG3 knockdown, possibly due to reduced sarcomeric density and increased disarray. HSPB7 and DNAJB6 knockdowns displayed high sarcomere scores (similar to Scramble control). For all images, scale bar = 100 *µ*M. Magenta: BAG3; Green: MYBPC3; Cyan: DAPI.

**Figure S9.**
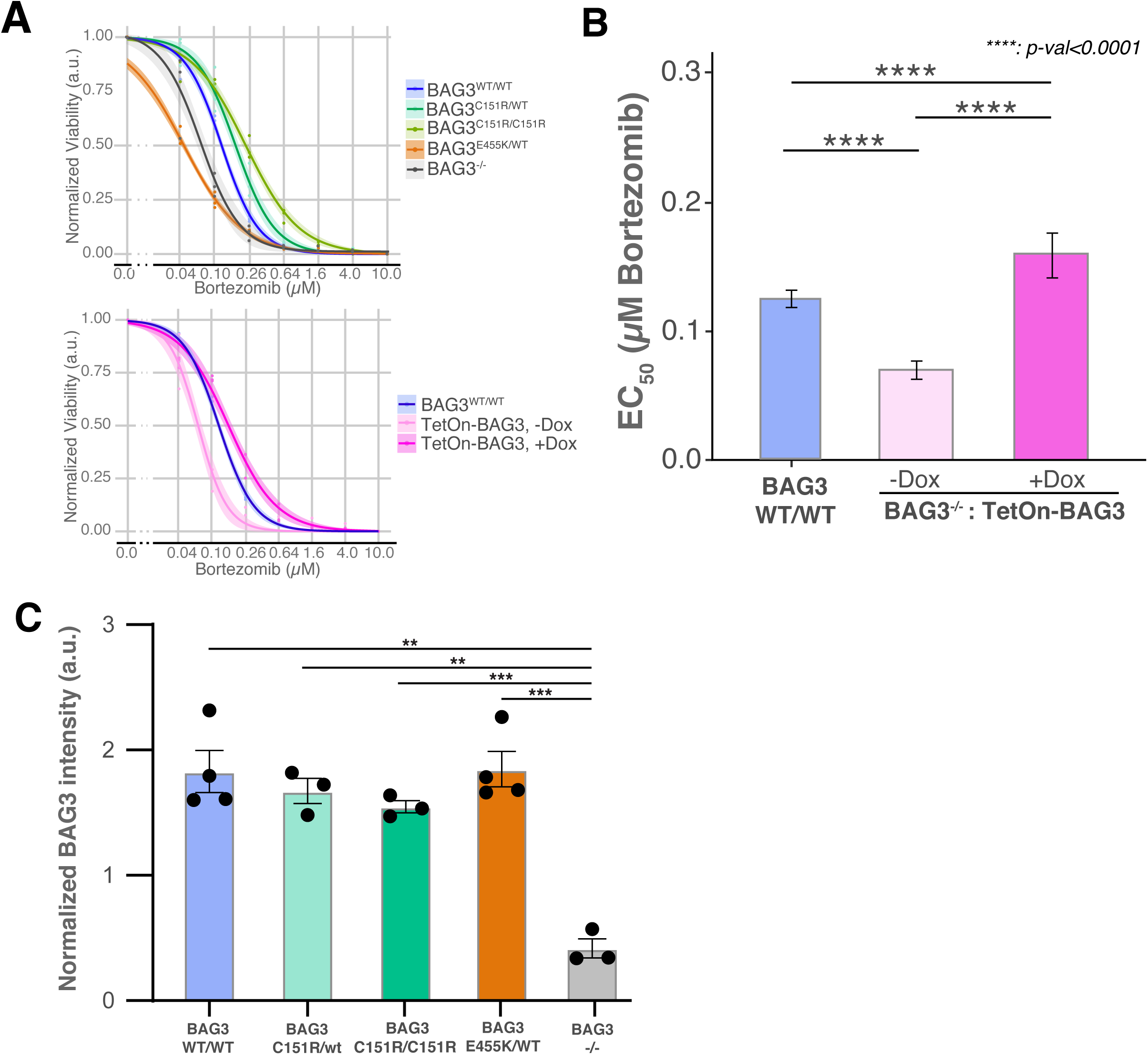
BAG3 overexpression rescues bortezomib sensitivity phenotype in BAG3^-/-^ cells, and BAG3^C151R^ and BAG3^E455K^ do not change BAG3 protein levels in iPS-CMs. (A) Bortezomib dose-response curves for the data used for EC_50_ calculations. (B) Calculated EC_50_ and 95% confidence intervals for Bortezomib in control (WT/WT), and BAG3^-/-^ iPS-CM with and without BAG3 overexpression. *N=*3. ****: P-value<0.0001 and *: p-value<0.5 using one-way ANOVA with post-hoc Zidak correction. (C) Capillary immunoassay (Simple Western) quantification of BAG3 protein levels in iPS-CM differentiated from iPSCs heterozygous or homozygous for the indicated BAG3 alleles. *N*=3. **: p-value<0.01; ***: p-value<0.001. One-way ANOVA with post-hoc Tukey multiple comparisons test.

