## Supplementary Figures for "Functional analysis of a common BAG3 allele associated with protection from heart failure"

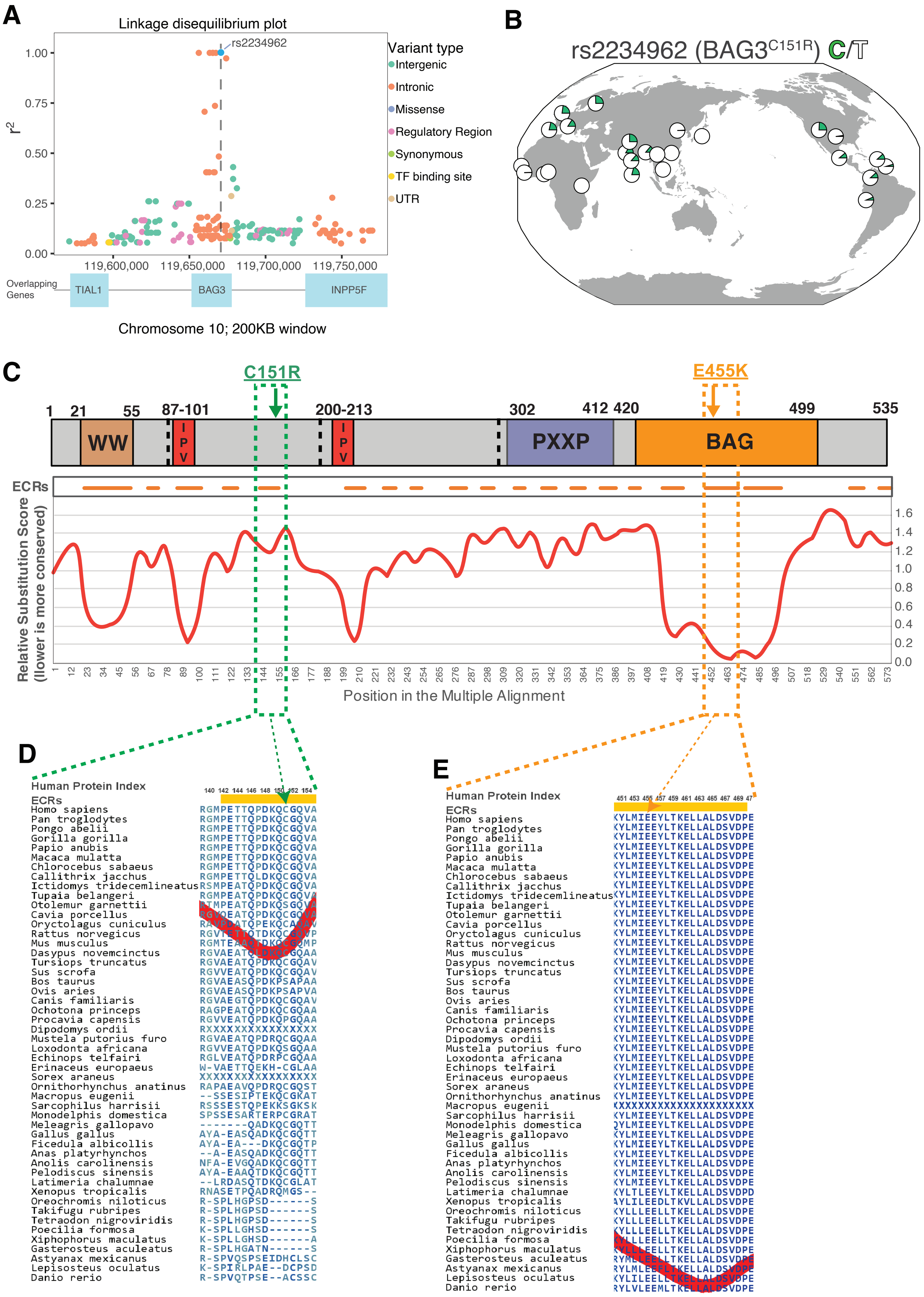
A
Linkage disequilibrium plot
rs2234962
Variant type
Intergenic
Intronic
Missense
Regulatory Region
Synonymous
TF binding site
UTR
r2
1.00
0.75
0.50
0.25
0.00
119,600,000
119,650,000
119,700,000
119,750,000
Overlapping Genes
TIAL1
BAG3
INPP5F
Chromosome 10; 200KB window
B
rs2234962 (BAG3C151R) C/T
C
C151R
E455K
1 21 55 87-101 200-213 302 412 420 499 535
WW
IPV
IPV
PXXP
BAG
ECRs
Relative Substitution Score (lower is more conserved)
1.6
1.4
1.2
1.0
0.8
0.6
0.4
0.2
0.0
Position in the Multiple Alignment
D
Human Protein Index
140 142 144 146 148 150 152 154
ECRs
Homo sapiens RGMPETTQPDKQCGQVA
Pan troglodytes RGMPETTQPDKQCGQVA
Pongo abelii RGMPETTQPDKQCGQVA
Gorilla gorilla RGMPETTQPDKQCGQVA
Papio anubis RGMPETTQPDKQCGQVA
Macaca mulatta RGMPETTQPDKQCGQVA
Chlorocebus sabaeus RGMPETTQPDKQCGQVA
Callithrix jacchus RGMPETTQLDKQCGQVA
Ictidomys tridecemlineatus RSMPEATQPDKQCGQVA
Tupaia belangeri
Otolemur garnettii
Cavia porcellus
Oryctolagus cuniculus
Rattus norvegicus
Mus musculus
Dasypus novemcinctus
Tursiops truncatus RGVAEATQPDKQCGQAV
Sus scrofa RGVVEATQPDKQCGQAV
Bos taurus RGVAEASQPDKPSAPAA
Ovis aries RGVAEASQPDKPSAPVA
Canis familiaris RGVAEGTQPDKQCGQAV
Ochotona princeps RAGPEATQPDKQCGQAA
Procavia capensis RGVVEATQPDKQPGQAA
Dipodomys ordii RXXXXXXXXXXXXXXXX
Mustela putorius furo RGVAEATQPDRQCGQAA
Loxodonta africana RGVVEATQPDKQSGQAA
Echinops telfairi RGLVEATQPDRPCGQAA
Erinaceus europaeus W-VAETTQEKH-CGLAA
Sorex araneus XXXXXXXXXXXXXXXXX
Ornithorhynchus anatinus RAPAEAVQPDRQCGQST
Macropus eugenii --SSESIPTEKQCGKAT
Sarcophilus harrisii RSSSESTQPEKKSGKSK
Monodelphis domestica SPSSESARTERPCGRAT
Meleagris gallopavo -------QADKQCGQTT
Gallus gallus AYA-EASQADKQCGQTT
Ficedula albicollis AYA-EA---DKQCGQTP
Anas platyrhynchos --A-EASQADKQCGQTT
Anolis carolinensis NFA-EVGQADKQCGQTT
Pelodiscus sinensis AYA-EAAQTDKQCGQTT
Latimeria chalumnae --LRDASQTDKQCGLAT
Xenopus tropicalis RNASETPQADRQMGS--
Oreochromis niloticus R-SPLHGPSD------S
Takifugu rubripes R-SPLHGPSD------S
Tetraodon nigroviridis R-SPLHGPSD------S
Poecilia formosa K-SPLLGHSD------S
Xiphophorus maculatus K-SPLLGHSD------A
Gasterosteus aculeatus R-SPLHGATN------S
Astyanax mexicanus R-SPVQSPSEIDHCLSC
Lepisosteus oculatus K-SPIRLPAE--DCPSD
Danio rerio R-SPVQTPSE--ACSSC
E
Human Protein Index
451 453 455 457 459 461 463 465 467 469 47
ECRs
Homo sapiens KYLMIEEYLTKELLALDSVDPE
Pan troglodytes KYLMIEEYLTKELLALDSVDPE
Pongo abelii KYLMIEEYLTKELLALDSVDPE
Gorilla gorilla KYLMIEEYLTKELLALDSVDPE
Papio anubis KYLMIEEYLTKELLALDSVDPE
Macaca mulatta KYLMIEEYLTKELLALDSVDPE
Chlorocebus sabaeus KYLMIEEYLTKELLALDSVDPE
Callithrix jacchus KYLMIEEYLTKELLALDSVDPE
Ictidomys tridecemlineatus KYLMIEEYLTKELLALDSVDPE
Tupaia belangeri KYLMIEEYLTKELLALDSVDPE
Otolemur garnettii KYLMIEEYLTKELLALDSVDPE
Cavia porcellus KYLMIEEYLTKELLALDSVDPE
Oryctolagus cuniculus KYLMIEEYLTKELLALDSVDPE
Rattus norvegicus KYLMIEEYLTKELLALDSVDPE
Mus musculus KYLMIEEYLTKELLALDSVDPE
Dasypus novemcinctus KYLMIEEYLTKELLALDSVDPE
Tursiops truncatus KYLMIEEYLTKELLALDSVDPE
Sus scrofa KYLMIEEYLTKELLALDSVDPE
Bos taurus KYLMIEEYLTKELLALDSVDPE
Ovis aries KYLMIEEYLTKELLALDSVDPE
Canis familiaris KYLMIEEYLTKELLALDSVDPE
Ochotona princeps KYLMIEEYLTKELLALDSVDPE
Procavia capensis KYLMIEEYLTKELLALDSVDPE
Dipodomys ordii KYLMIEEYLTKELLALDSVDPE
Mustela putorius furo KYLMIEEYLTKELLALDSVDPE
Loxodonta africana KYLMIEEYLTKELLALDSVDPE
Echinops telfairi KYLMIEEYLTKELLALDSVDPE
Erinaceus europaeus KYLMIEEYLTKELLALDSVDPE
Sorex araneus KYLMIEEYLTKELLALDSVDPE
Ornithorhynchus anatinus KYLMIEEYLTKELLALDSVDPE
Macropus eugenii XXXXXXXXXXXXXXXXXXXXXX
Sarcophilus harrisii KYLMIEEYLTKELLALDSVDPE
Monodelphis domestica QYLMIEEYLTKELLALDSVDPE
Meleagris gallopavo KYLMIEEYLTKELLALDSVDPE
Gallus gallus KYLMIEEYLTKELLALDSVDPE
Ficedula albicollis KYLMIEEYLTKELLALDSVDPE
Anas platyrhynchos KYLMIEEYLTKELLALDSVDPE
Anolis carolinensis KYLMIEEYLTKELLALDSVDPE
Pelodiscus sinensis KYLMIEEYLTKELLALDSVDPE
Latimeria chalumnae KYLTLEEYLTKELLALDSVDPD
Xenopus tropicalis AYLILEEDLTKVLLALDSVDPE
Oreochromis niloticus KYLLLEELLTKELLALDSVDPE
Takifugu rubripes KYLLLEELLTKELLALDSVDPE
Tetraodon nigroviridis KYLLLEELLTKELLALDSVDPE
Poecilia formosa
Xiphophorus maculatus
Gasterosteus aculeatus
Astyanax mexicanus
Lepisosteus oculatus
Danio rerio

**Figure S1. Expanded genetic data graphs for rs2234962, BAG3$^{C151R}$. Conservation of residues affected by BAG3$^{C151R}$ and BAG3$^{E455K}$ variants. (A)** Zoomed out version of Figure 1A, showing a window of 200KB. Dot color indicates type of nucleotide change. **(B)** Allele frequency map for rs2234962 depicting all 1000 Genomes populations. **(C)** Top: Diagram of BAG3 domain structure. Bottom: Amino acid conservation plot for matching BAG3 regions. Decreasing Relative Substitution Score regions (valleys) indicate sequences with high conservation across species and are annotated as Evolutionary Constrained Regions (ECRs). (D-E) Zoomed in regions for the ECRs around BAG3$^{C151}$(D) and BAG3$^{E455}$(E).

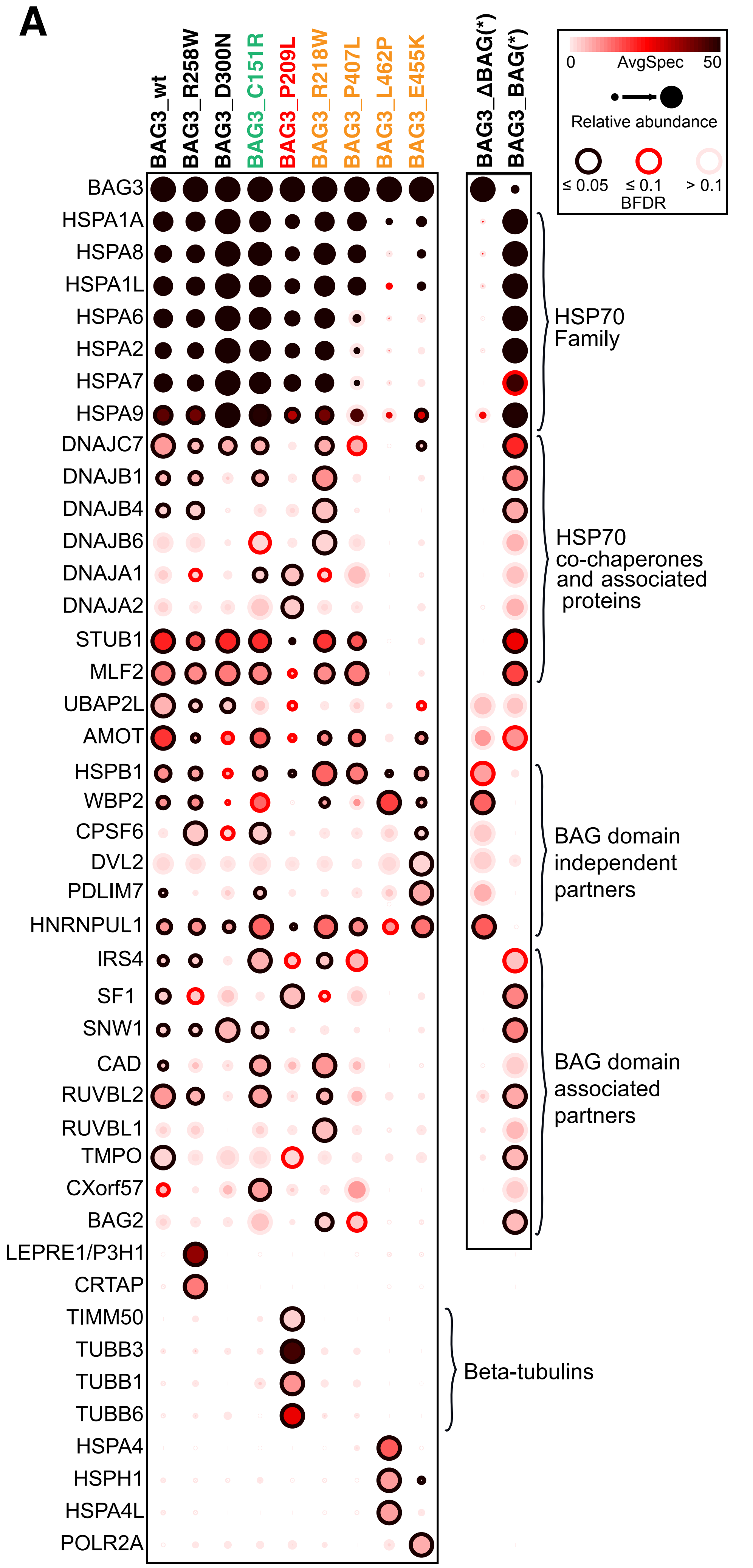


**Figure S2. Co-precipitation profiles of different BAG3 variants overexpressed in a HEK293 cell background.** Dot size represents the amount of co-precipitated protein normalized across variants. Dot color represents absolute protein abundance (spectral counts). Dot rim represents statistical significance. Yellow colored variants are known pathogenic variants associated with DCM. Green variant is putative cardioprotective variant $BAG3^{C151R}$. Red variant ($BAG3^{P209L}$) is associated with skeletal myofibrillar myopathy. Black variants are not associated to DCM or any other pathology. Rightmost two columns depict data for BAG3 truncated variant without the BAG domain (BAG3_ ΔBAG) and for the BAG3 protein BAG domain only (BAG3_ΔBAG). For the truncated variants, no normalization by bait levels was performed. *N*=5, BFDR obtained using SAINTexpress (see methods).

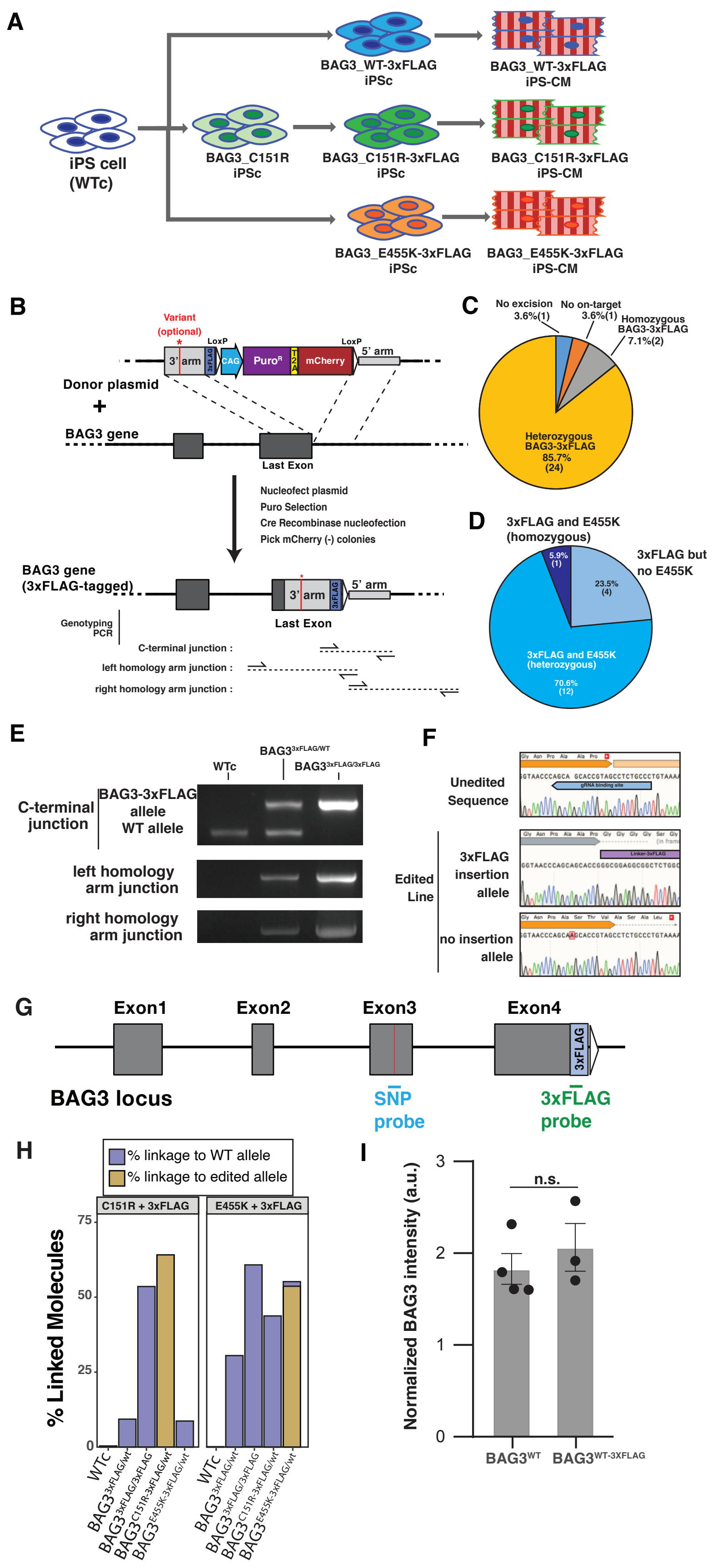


**Figure S3. Generation of the isogenic cell lines carrying BAG3 variants and a 3xFLAG epitope tag fusion in the endogenous copy of the BAG3 gene.** (A) Workflow for the cell line generation. (B) Strategy for the insertion of a 3xFLAG epitope fusion at the C-terminal of the BAG3 gene. The BAG3$^{C151R\text{-}FLAG}$ variant was generated using the same process on a preexisting cell line bearing the C151R mutation. To generate the BAG3$^{E455K\text{-}FLAG}$ cell line, the homology arms were engineered to contain the SNP and insert it during recombination. (C-D) Genotypes of the single-cell clones picked for 3xFLAG insertion (C) and the co-segregation of the BAG3$^{E455K}$ variant (D). (E) Genotyping the products of the 3xFLAG insertion by PCR. (F) Cells with a heterozygous insertion of the 3xFLAG epitope tag also had a SNP in the other allele that extended the BAG3 protein product by 4 amino acids. (G-H) A droplet digital PCR phasing test was used to select clones that contained the desired SNP variants and the 3xFLAG C-terminal sequence in the same allele. The test used different probes (G) to generate an estimate of linked molecules for each cell line and probe combination (H; See Methods for more details) (I) Insertion of the 3xFLAG fusion in the BAG3 gene did not alter the protein levels. *N*=3; one-way ANOVA.

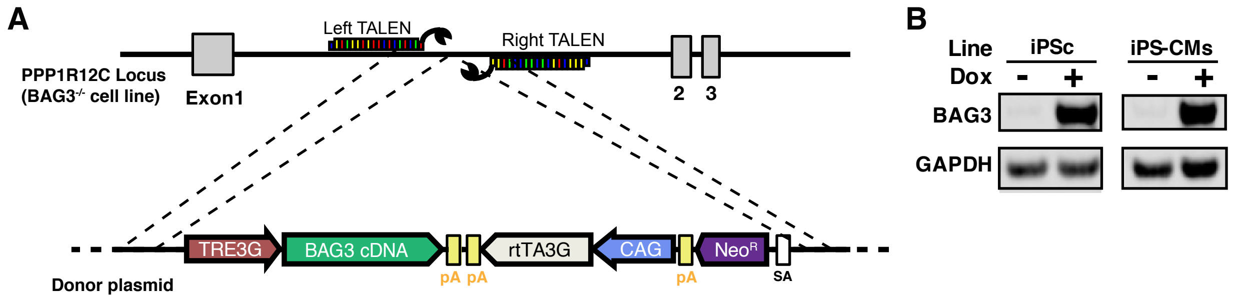


**Figure S4. Generation of a cell line with inducible expression of the BAG3$^{WT}$ protein.** (A) Diagram of the editing strategy. On a BAG3$^{-/-}$ cell background, a doxycycline-activated BAG3-3xFLAG expression cassette was inserted in the PPP1R12C (AAVS1) safe-harbor locus. (B) Western blot of the BAG3 expression on BAG3$^{-/-}$:TetOn-BAG3$^{WT\text{-}3xFLAG}$ iPSCs and iPS-CMs with and without Doxycycline addition.

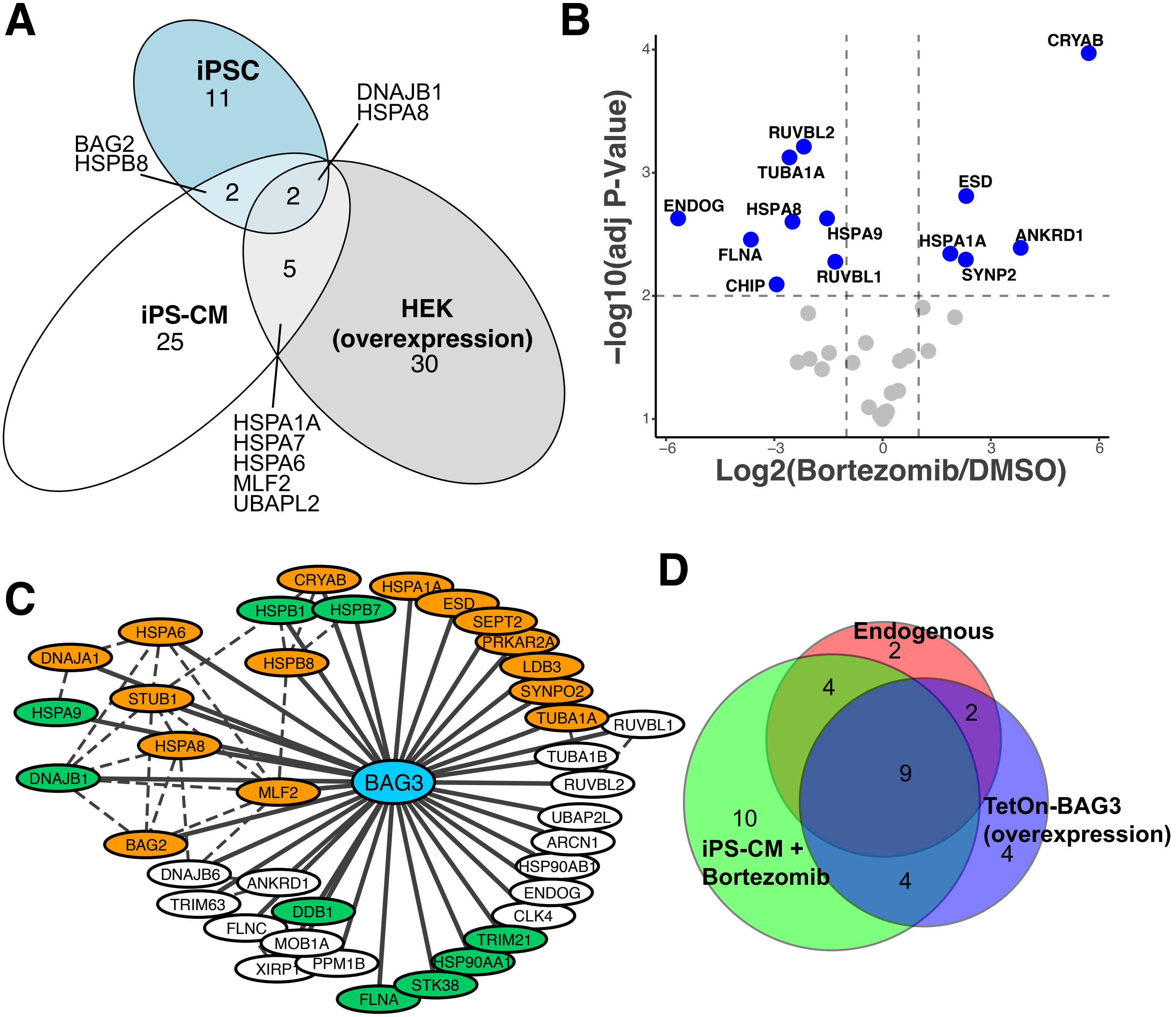


**Figure S5. Affinity purification - mass spectrometry characterization of BAG3 binding partners in a cardiomyocyte background.** (A) Venn diagram of the high confidence BAG3$^{WT}$ protein-protein interactions identified in three cellular backgrounds. HEK293T cells had overexpressed baits, while iPSC have much lower levels of endogenous BAG3 expression than iPS-CM, which could have influenced the results. Each cell type specific dataset was scored separately against its own matched control(s). (B) Volcano plots depicting co-precipitation intensity in BAG3$^{WT}$ cardiomyocytes treated with Bortezomib (100nM) relative to DMSO (1:10.000) for 24hours. Horizontal dashed line indicates statistical significance threshold (adjusted p-value <0.01) and vertical dashed lines indicate a fold change of 2. *N*=4. (C) Network diagram of the iPS-CM co-precipitation partners identified for BAG3 in this study. Nodes in orange indicate partners that significantly changed when pulling down BAG3$^{E455K}$. Nodes in Green indicate partners that significantly changed when pulling down BAG3$^{C151R}$. Dashed lines: known interactions in the iRefIndex database. (D) Venn diagram comparing the BAG3 binding partners identified in an iPS-CM background when using endogenous basal levels of expression, an overexpression system, or endogenous expression under proteotoxic stress. The graph highlights the importance of using endogenous expression for accurate characterization of binding partners, and the information gained from using a stress state.

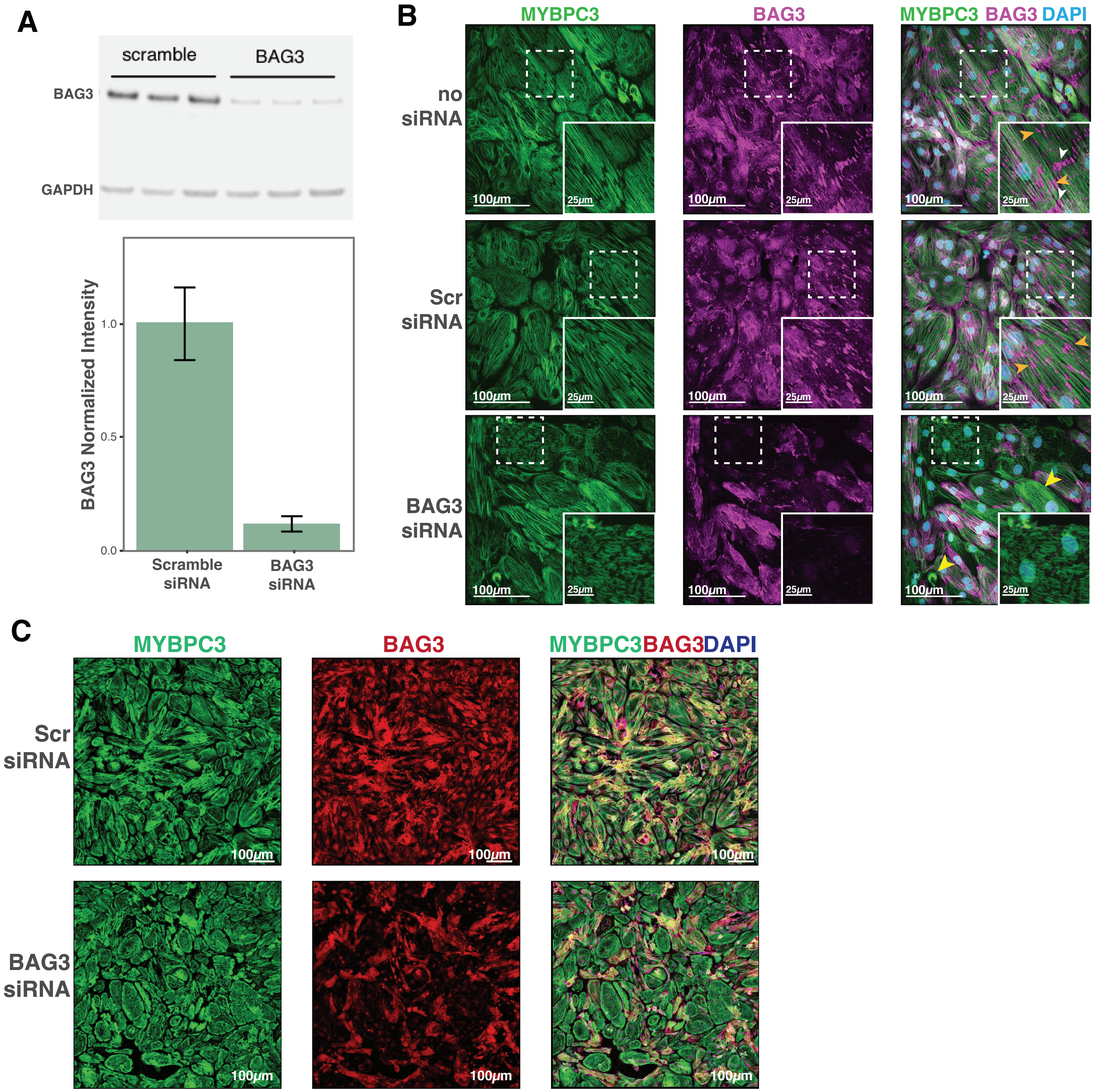


**Figure S6. Additional sample micrographs from BAG3 knockdown iPS-CM.** (A) BAG3 silencing by siRNA was effective at reducing protein levels (~85% reduction). *N*=3. (B) Additional sample micrographs from BAG3- and Scr-siRNA-treated iPS-CM, plus a no-siRNA condition. Orange arrowheads: BAG3 accumulation on myofibrillar breaks; white arrowheads: BAG3 accumulation on polar ends of cells; yellow arrowheads: iPS-CM displaying myofibrillar aggregation and collapse. (C) Sample images in the same magnification used for the automated scoring analysis. Lower magnification allowed for faster acquisition and richer features to use directly in the scoring scheme, but higher magnification images were used elsewhere in this manuscript for easier viewing.

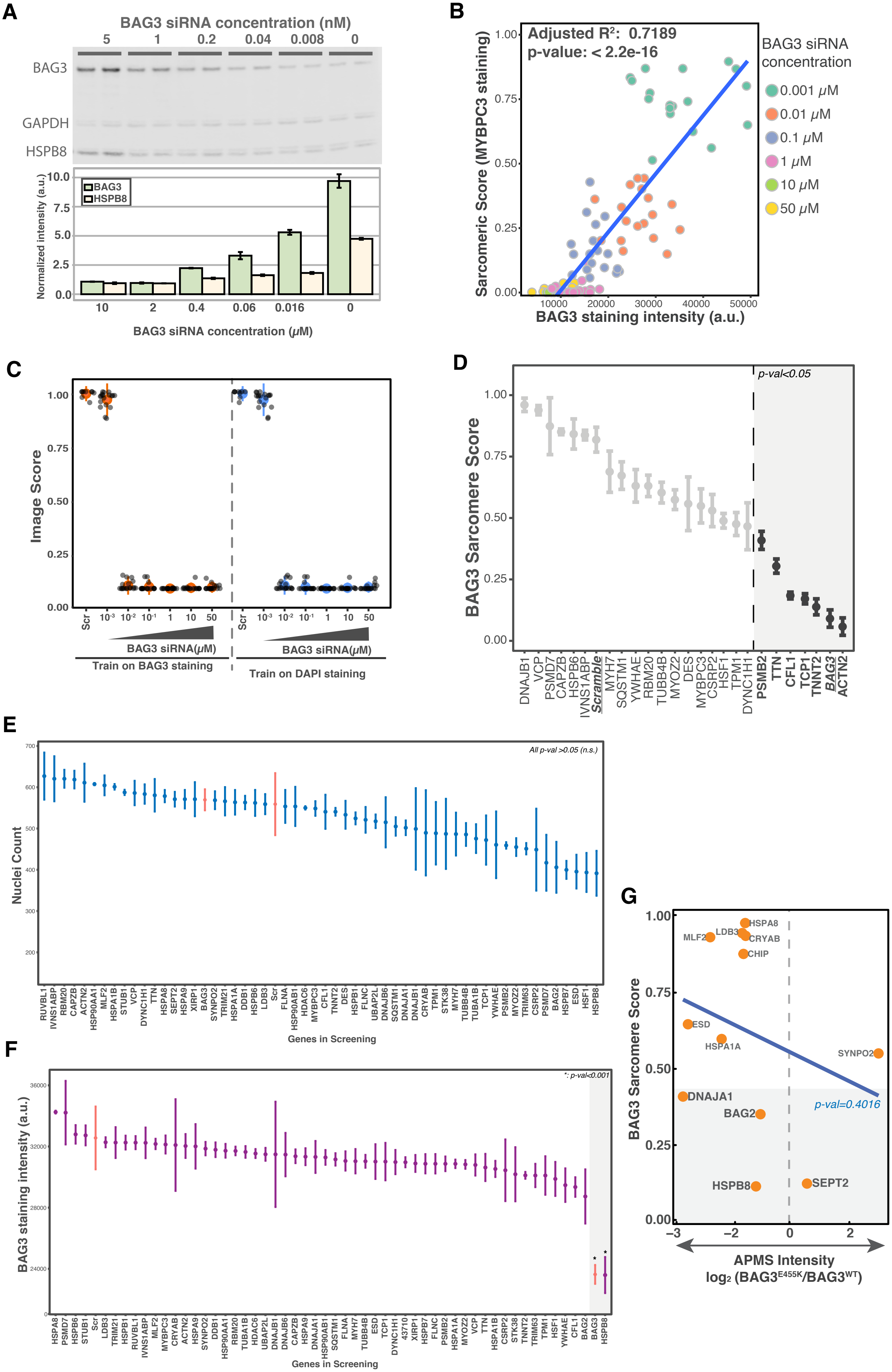
A
BAG3 siRNA concentration (nM)
5
1
0.2
0.04
0.008
0
BAG3
GAPDH
HSPB8
Normalized intensity (a.u.)
BAG3
HSPB8
10
2
0.4
0.06
0.016
0
BAG3 siRNA concentration (μM)
B
Adjusted R²: 0.7189
p-value: < 2.2e-16
Sarcomeric Score (MYBPC3 staining)
BAG3 staining intensity (a.u.)
BAG3 siRNA concentration
0.001 μM
0.01 μM
0.1 μM
1 μM
10 μM
50 μM
C
Image Score
Scr
BAG3 siRNA(μM)
Train on BAG3 staining
Train on DAPI staining
D
p-val<0.05
BAG3 Sarcomere Score
DNAJB1
VCP
PSMD7
CAPZB
HSPB6
IVNS1ABP
Scramble
MYH7
SQSTM1
YWHAE
RBM20
TUBB4B
MYOZ2
DES
MYBPC3
CSRP2
HSF1
TPM1
DYNC1H1
PSMB2
TTN
CFL1
TCP1
TNNT2
BAG3
ACTN2
E
All p-val >0.05 (n.s.)
Nuclei Count
Genes in Screening
F
*: p-val<0.001
BAG3 staining intensity (a.u.)
Genes in Screening
G
BAG3 Sarcomere Score
HSPA8
LDB3
MLF2
CRYAB
CHIP
ESD
HSPA1A
SYNPO2
DNAJA1
BAG2
p-val=0.4016
HSPB8
SEPT2
APMS Intensity
log₂ (BAG3^E455K/BAG3^WT)

**Figure S7. Quality control and additional data from the siRNA knockdown-myofibrillar scoring workflow.** (A) Western blot demonstrating the titration of BAG3 siRNA results in decreasing cellular BAG3 protein levels. Shown at the bottom of the western blot are levels of HSPB8, which are also affected by BAG3 knockdown. *N=2.* (B) BAG3 sarcomere score inversely correlates with BAG3 protein levels. *N* = 18 images. (C) Plot of the image scores that result from training based on BAG3 or DAPI staining. These stains do not display the same dynamic range as scoring based on myofibrillar (MYBPC3) staining. *N*=9 for scramble; 18 for the rest. (D) BAG3 Sarcomere Score for the knockdown of selected factors that were not identified in our AP-MS coprecipitation studies. Dots represent mean of 3 replicates from separate wells, each being the median score of 9 images from the same well. Error bars: SEM. P-val cutoff: 0.05 using a one-way ANOVA with post-hoc Dunnett test. (E-F) Plot of the nuclei count(E) and BAG3 staining(F) intensities for the gene knockdowns used in the siRNA-myofibrillar scoring analyses. Dots represent mean of 3 replicates from separate wells, each being the median score of 9 images from the same well. Error bars: SEM. P-val cutoff: 0.05 using a one-way ANOVA with post-hoc Dunnett test. (G) Plotting of the APMS intensity ratio for $BAG3^{E455K}$ differential interactors and their BAG3 Sarcomere Score. There is no statistically significant correlation. P-value obtained fitting a linear model. Pearson's product-moment correlation: -0.27.

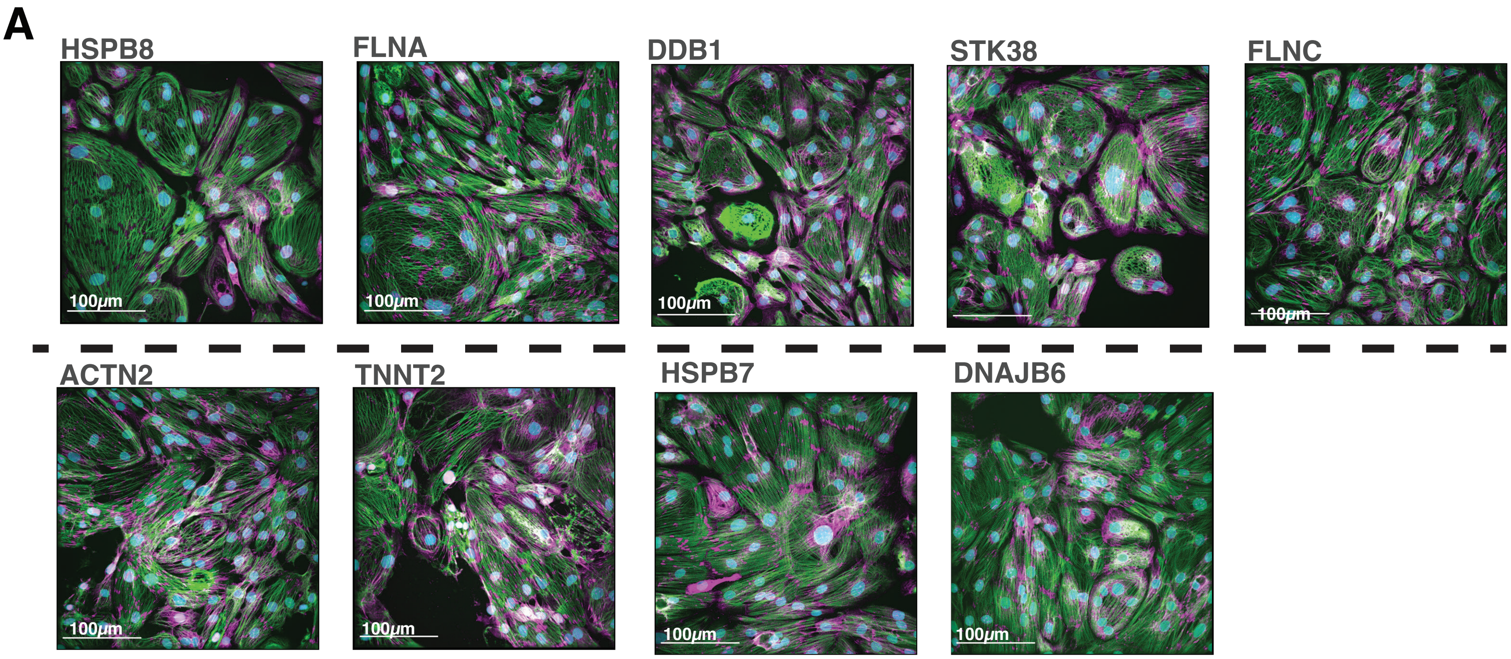


**Figure S8. Sample images from selected siRNA knockdowns.** HSPB8 knockdown was the only knockdown to significantly reduce BAG3 levels. FLNA, DDB1 and STK38 are $BAG3^{C151R}$ differential interactors whose knockdown resulted in sarcomere scores similar to BAG3 knockdown. ACTN2 and TNNT2 are well known sarcomere components that display low sarcomere scores similar to BAG3 knockdown, possibly due to reduced sarcomeric density and increased disarray. HSPB7 and DNAJB6 knockdowns displayed high sarcomere scores (similar to Scramble control). For all images, scale bar = 100 $\mu$M. Magenta: BAG3; Green: MYBPC3; Cyan: DAPI.

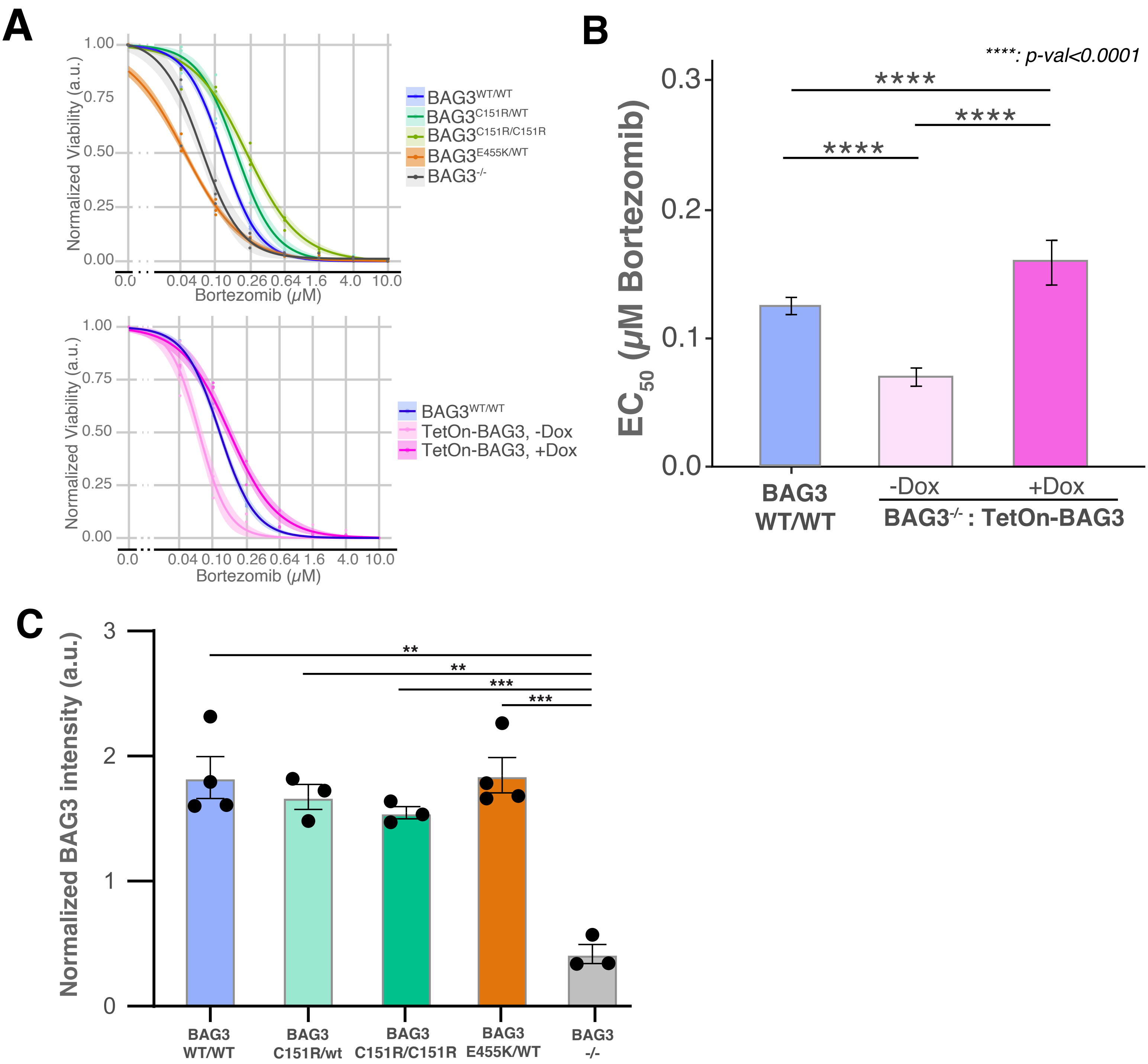


**Figure S9. BAG3 overexpression rescues bortezomib sensitivity phenotype in BAG3$^{-/-}$ cells, and BAG3$^{C151R}$ and BAG3$^{E455K}$ do not change BAG3 protein levels in iPS-CMs.** (A) Bortezomib dose-response curves for the data used for $EC_{50}$ calculations. (B) Calculated $EC_{50}$ and 95% confidence intervals for Bortezomib in control (WT/WT), and BAG3$^{-/-}$ iPS-CM with and without BAG3 overexpression. *N*=3. ****: P-value<0.0001 and *: p-value<0.5 using one-way ANOVA with post-hoc Zidak correction. (C) Capillary immunoassay (Simple Western) quantification of BAG3 protein levels in iPS-CM differentiated from iPSCs heterozygous or homozygous for the indicated BAG3 alleles. *N*=3. **: p-value<0.01; ***: p-value<0.001. One-way ANOVA with post-hoc Tukey multiple comparisons test.
